# Influence of expression and purification protocols on Gα biochemical activity: kinetics of plant and mammalian G protein cycles

**DOI:** 10.1101/2023.05.10.540258

**Authors:** Timothy E. Gookin, David Chakravorty, Sarah M. Assmann

## Abstract

Heterotrimeric G proteins are a class of signal transduction complexes with broad roles in human health and agriculturally important plant traits. In the classic paradigm, guanine nucleotide binding to the Gα subunit regulates the activation status of the complex. Using the *Arabidopsis thaliana* Gα subunit, GPA1, we developed a rapid StrepII-tag mediated purification method that facilitates isolation of protein with increased enzymatic activities as compared to conventional methods, and is demonstrably also applicable to mammalian Gα subunits. We subsequently utilized domain swaps of GPA1 and human GNAO1 to demonstrate the instability of recombinant GPA1 is a function of the interaction between the Ras and helical domains, and can be partially uncoupled from the rapid nucleotide binding kinetics displayed by GPA1.

## Introduction

The heterotrimeric G protein complex consists of an alpha (Gα), beta (Gβ), and gamma (Gγ) subunit, in which Gβ and Gγ exist as a non-dissociable dimer. Heterotrimeric G proteins (G proteins) are well-studied conserved eukaryotic signal transduction components. Mutations of G protein subunits in humans have been associated with diseases and developmental abnormalities including cancer [1–3], neurodevelopmental disorders [4], McCune-Albright Syndrome [5], diabetes [6,7], hypertension [8] and ventricular tachycardia [9]. In plants, null mutants of G protein subunits have been utilized to implicate G proteins in agronomically important traits such as morphological development [10,11], grain shape and yield [12], hormone sensitivity [13,14], stomatal responses [15–17], salinity tolerance [18], drought tolerance [19] and pathogen resistance [20,21].

The Gα subunit of the G protein heterotrimer binds guanine nucleotides in a binding pocket located within a cleft between the Ras-like and helical domains of the protein. The identity of the nucleotide, GDP or GTP, determines the activation state of the heterotrimer in the canonical signaling paradigm. In the inactive heterotrimer, Gα exists in the GDP-bound form, while exchange of GDP for GTP results in activation of the heterotrimer and dissociation of Gα from the Gβγ dimer. Gα and Gβγ are then able to signal to downstream effectors, until the intrinsic GTPase activity of the Gα subunit hydrolyzes GTP to GDP, thereby stimulating reassociation of the inactive heterotrimer. The activation status of the complex can be regulated by guanine nucleotide exchange factors (GEFs) including 7-transmembrane spanning G protein-coupled receptors (GPCRs) that stimulate GTP-binding, and GTPase activating proteins (GAPs) such as regulator of G protein signaling (RGS) proteins that stimulate GTP hydrolysis [22]. In mammals the GPCR superfamily is large and perceives diverse ligands, with over 800 GPCRs encoded in the human genome [23]. In contrast, only a few 7TM proteins have been identified as candidate GPCRs in plants [24,25], including in the intensively studied model dicot *Arabidopsis thaliana* [26]. Receptor-like kinases (RLKs) may predominate in the GPCR role in plants [17,27–31].

Purification of active recombinant heterotrimeric G protein Gα subunits is integral to understanding structure-function relationships. Purification of full-length functional Gα subunits from *E. coli* has presented challenges for some Gα family members; for example, Gα_t_ consistently aggregates within inclusion bodies [32]. Previously employed solutions for recombinant Gα expression have included N-terminal protein deletions [33,34], chimeric substitutions [35–37] and use of insect cell expression systems [38,39]. Here, we sought to assess the enzyme kinetics of the sole canonical Arabidopsis Gα subunit, GPA1, in comparison with two closely related mammalian G proteins in the Gα_i_ family, GNAI1 and GNAO1. In the course of our investigations we addressed some of the challenges to purifying Gα proteins, finding that specific purification protocols markedly affect the activity of both plant and mammalian Gα proteins.

GPA1 has previously been described to: i) display self-activating properties due to spontaneous nucleotide exchange and fast GTP-binding, and ii) exhibit slow GTPase activity, which skews the protein to the GTP-bound form, especially when compared to the human Gα_i1_ protein, GNAI1 [33,40]. Jones et al. [33] determined the structure of GPA1 by x-ray crystallography, discovering that the GPA1 tertiary structure bears a strong resemblance to that of GNAI1. Yet, GPA1 and GNAI1 also display distinctly different enzymatic activities, indicating that differences arise on a finer scale, possibly due to high levels of intrinsic disorder within, and dynamic motion of, the GPA1 helical domain [33]. To further investigate Gα enzymatic properties, we developed a robust expression and rapid purification protocol utilizing dual StrepII-tags [41], which allowed Gα elution in a buffer directly compatible with downstream nucleotide binding and GTP hydrolysis assays, thereby abrogating any need for protracted buffer component removal, e.g. by dialysis. GPA1 exhibits increased activity when rapidly purified using our specific purification protocol. Furthermore, assays of the two Gα subunits from the most closely related human Gα family, Gα_i_, demonstrate that mammalian Gα activity is also impacted by the choice of purification regime, and that our StrepII-tag approach to expression and purification is applicable to mammalian Gα subunits. Our results also imply that the StrepII-tag approach may represent an improved protocol for purification of other classes of enzymes including but not limited to small G proteins and other nucleotidases, due to the direct compatibility of the elution buffer with downstream applications.

## Materials and methods

### Cloning

*GPA1* was amplified from *Arabidopsis* cDNA with flanking NcoI and BspEI restriction sites. *GNAI1* with the same flanking restriction sites was amplified from a wild type clone (Genscript, clone OHu13586) and from a designed codon harmonized [42] gBlock synthesized by Integrated DNA Technologies. These Gα subunits were cloned into the NcoI and BspEI sites of pSTTa, a vector we adapted from pGEX to include N-terminal dual-StrepII tags, thrombin and TEV protease sites, a multiple cloning site and an optional C-terminal FLAG tag. *GNAO1* was amplified from a commercial clone (Genscript, clone OHu15183), adapting a 5’ BspHI restriction site (yields a sticky end compatible with NcoI) and a blunt 3’ end to clone into NcoI/PmlI sites of pSTTa. The C-terminal RGS box of *RGS1* [43], corresponding to residues 247-459, was amplified from *Arabidopsis* cDNA with flanking NcoI and BspEI sites to clone into pSTTa in the same manner as *GPA1* and *GNAI1*. All cDNAs cloned into pSTTa included a stop codon, so the open reading frame (ORF) did not read through to the C-terminal FLAG tag included in the vector. Mutants of *GPA1*, *GNAI1* and *GNAO1* were generated by REPLACR mutagenesis [44]. *GPA1-GNAO1* helical domain swaps were generated by overlap-extension PCR [45] and cloned into pSTTa as above, with the exception that the *GPA1^GNAO1hel^* construct was amplified with a 5’ BspHI site. Helical domains were defined as GPA1 residues E68-Y188 and GNAO1 residues G63-R177, with the flanking sequences, GPA1 residues M1-D67 and A189-L383 or GNAO1 residues M1-S62 and T178-Y354, defined as the Ras domains, consistent with the regions used in the GPA1-GNAI1 domain swap performed by Jones et al. [33]. His- and GST-tagged constructs were generated by amplifying the ORFs of *GPA1*, *GNAI1* and *GNAO1*, which were A-tailed, TOPO cloned into pCR8 and mobilized by LR Gateway recombination into pDEST17 (for His-tagged expression), and in the case of *GPA1*, pDEST15 (for GST-tagged expression comparisons) (Thermo Fisher Scientific). Primers for ORF cloning, mutagenesis and overlap-extension PCRs are listed in Table S1. All sequences were verified by Sanger sequencing.

### Protein expression

Proteins were heterologously expressed in *E. coli* BL21 DE3 cells using 75 μg/ml carbenicillin for plasmid selection. Typically, fresh transformants were grown in 7.5 ml overnight cultures (LB media supplemented with 0.5% D-glucose (w/v) and 3 g/L MgCl-_2_), pelleted by centrifugation at 5000 g for 10 minutes, resuspended in 5 ml fresh pre-warmed LB, and grown at 37°C. Five ml of pre-warmed HSLB (LB media supplemented with 17 g/L NaCl and 3 g/L MgCl-_2_, pH 7.0) was added at T=20 and 40 minutes. At T=60 minutes the pre-culture was added to 600 ml prewarmed HSLB in a vigorously shaking (225 rpm) 2 L baffled flask (OD_600_ = 0.04-0.06). Cultures were grown to an OD_600_ of 0.7-0.8, transferred to a room temperature (20-21 °C) shaker and grown for 20 minutes before induction with 125 μM IPTG for 3-4 hours. Cells were pelleted by 6000 g centrifugation for 10 minutes at 4 °C. Cell pellets were promptly frozen at −80 °C and typically processed the following morning, though proteins retained activity when cell pellets were stored for multiple weeks at −80 °C.

### Protein purification

All buffers were prepared with high purity premium grade reagents (e.g. Honeywell TraceSelect, Millipore Sigma BioXtra or EMD Millipore EmSure) to minimize introduction of extraneous metals, and supplemented with one tablet Complete EDTA-free protease inhibitor (Roche, 5056489001) or Pierce Protease Inhibitor Tablets, EDTA-free (Thermo Fisher Scientific, A32965) per 50 ml. Columns were pre-rinsed with 1 ml of 0.25% Tween-20. Frozen cell pellets containing expressed StrepII-tag fusion proteins were resuspended with a 10 ml Pasteur pipet in 10 ml buffer W1 (100 mM Tris-HCl, 500 mM NaCl, 2 mM MgCl_2_, 5 mM TCEP and 5% glycerol pH 8.0) supplemented with ∼10 mg lysozyme (Millipore Sigma, L1667), 25 μl/ml BioLock biotin blocker (IBA, 2-0205-050) and 5 μl Pierce Universal Nuclease (Thermo Fisher Scientific, 88702), and kept on ice. Cells were lysed by three rounds of sonication on ice using a Fisher Sonic Dismembrator equipped with a 3 mm tip with 1 second on/off pulses set to 20% amplitude for 15 seconds (i.e. 15x one second pulses over a 30-second timespan), and the cell debris was pelleted by centrifugation at 10000 g at 4 °C for 10-20 minutes. The supernatant was passed through a 0.2 μm PES filter directly into a 1 ml column, with a 6 ml total capacity, containing a 0.25 ml resin bed of Streptactin sepharose (IBA, 2-1201-010) pre-washed with buffer W1. Loaded columns were washed sequentially with 0.5 ml W1 (1x) and 0.3 ml W2 (3x) (50 mM Tris-HCl, 100 mM NaCl and 5% glycerol pH 7.7) before eluting with sequential fractions of 220, 350, and 165 μl of “EB base” (25 mM Tris-HCl, 50 mM NaCl and 5% glycerol pH 7.4) supplemented with freshly added 5 mM desthiobiotin (Millipore Sigma, D1411) to form “EB”. The identification of the minor contaminate DnaK was performed via gel band excision, NH_4_HCO_3_/CH_3_CN destaining, dehydration, and subsequent MS/MS sequencing by the Penn State College of Medicine Mass Spectrometry and Proteomics Facility.

For GST-fusion proteins, cell pellets were resuspended in TBS-NoCa binding buffer (50 mM Tris-HCl, 150 mM NaCl, 1 mM MgOAc, 10 mM β-mercaptoethanol and 1 mM Imidazole, pH 8.0) and sonicated and centrifuged as above. The resultant supernatant was passed through a 0.2 μm PES filter into a 1 ml column with a 0.25 ml Pierce Glutathione Agarose (Pierce, 16100) resin bed, essentially mimicking the StrepII purification protocol. Sequential washes were performed with 2 ml (x1) and 1 ml (x2) TBS-NoCa before protein elution with sequential fractions of 220 μl (E1), 350 μl (E2), and 165 μl (E3) TBS-NoCa supplemented with 10 mM glutathione.

His-fusion proteins were purified essentially as previously described for BODIPY reactions [46]. Briefly, our purification protocol mimicked the StrepII protocol, with the following modifications: lysis/binding buffer was replaced with 15 ml of 50 mM Tris-HCl pH 8.0, 100 mM NaCl, 2 mM MgCl_2_, 0.2% C_12_E_10_, supplemented with 5 µl β-mercaptoethanol post-sonication, cell debris was pelleted by centrifugation at 30000 g for 15 minutes, a 125 μl Talon (Takara, 635501) resin bed was used, the resin bed was washed with 1 ml of 50 mM Tris-HCl pH 8.0, 500 mM NaCl, 2 mM MgCl2, 5 mM β-mercaptoethanol, 0.2% C_12_E_10_ and 10 mM imidazole, and elution was performed with 20 mM Tris-HCl (pH 8.0), 250 mM NaCl, 5 mM β-mercaptoethanol, 10% glycerol and 250 mM imidazole.

Peak elution fractions (second eluate fraction; E2) of GST and His tagged proteins were subjected to buffer exchange using Amicon Ultra 0.5 ml 10 kDa cutoff columns (Millipore Sigma, UFC501024) with five sequential rounds of concentration performed by centrifugation at 14000 g and 4 °C for approximately 10 minutes and dilution with “EB base” (5x, 5x, 5x, 5x, 2x) for a total dilution of 1250x.

Protein quality and quantity were evaluated immediately after elution by SDS-PAGE of 10-20 μl fractions with a 3-4 lane mass ladder of Fraction V bovine serum albumin (BSA) (e.g. 0.5, 1.0, 1.5, 2.0 μg/lane) followed by Gel-Code Blue (Thermo Fisher Scientific, 24592) staining. Gels were imaged in a ChemiDoc MP Imaging System (Bio-Rad) and band intensity calculated using ImageLab software (Bio-Rad). Biochemical assays were initiated on the fraction displaying peak yield (almost always E2) immediately after PAGE analysis, generally 2-3 hours post-elution, during which time proteins had been stored on ice in a 4 °C refrigerator. We note that, under routine conditions and if pre-quantification of exact yield is not critical, the StrepII-tag E2 purity and concentration is consistent enough to allow for immediate biochemical analysis, within minutes of elution.

### BODIPY assays

BODIPY-GTP (BODIPY FL GTP, #G12411) and BODIPY-GTPγS (BODIPY™ FL GTP-γ-S, #G22183) stocks were purchased from Thermo Fisher Scientific and diluted to 100 nM in Tris-HCl pH 7.4 immediately prior to use. BSA or buffer alone was used as a negative control as indicated in each assay. Proteins and MgCl_2_ were diluted to twice the final assay concentration, generally 200 nM (GPA1, GNAO1 or BSA) or 400 nM (GNAI1 or BSA) supplemented with 10 mM MgCl_2_ in “EB base” on ice, normally in a master mix sufficient to perform reactions in triplicate. 100 μl of each diluted protein was aliquoted to wells of a Costar 96 well plate (Corning, 3631 - black with clear flat bottom non-treated plate) and loaded into a Synergy Neo2 multimode reader (Biotek), or in Figure 1C, an Flx800 plate reader (Biotek), set at 25 °C. Pre-injection background readings were taken with monochromators set to 486/18 nm excitation and 525/20 nm emission with a gain setting within the range 90-100 (Synergy Neo2), or 485/20 nm excitation and 528/20 nm emission filters with the sensitivity set to 90 (Flx800). Reactions were initiated utilizing plate reader injectors to dispense 100 μl of BODIPY-GTP or BODIPY-GTPγS to each well (at a rate of 250 μl/sec), yielding a final assay concentration of 50 nM BODIPY-GTP/-GTPγS, 100 or 200 nM protein and 5 mM Mg^2+^ cofactor. Kinetics were normally monitored in “plate mode” for 30 minutes with a kinetic interval of 3-6 seconds (Synergy Neo2) or 25-30 seconds (Flx800). For rapid monitoring of initial BODIPY-GTPγS binding rates, samples were monitored in “well mode” for 30 seconds with an 80 msec kinetic interval (Synergy Neo2).

**Figure 1.**
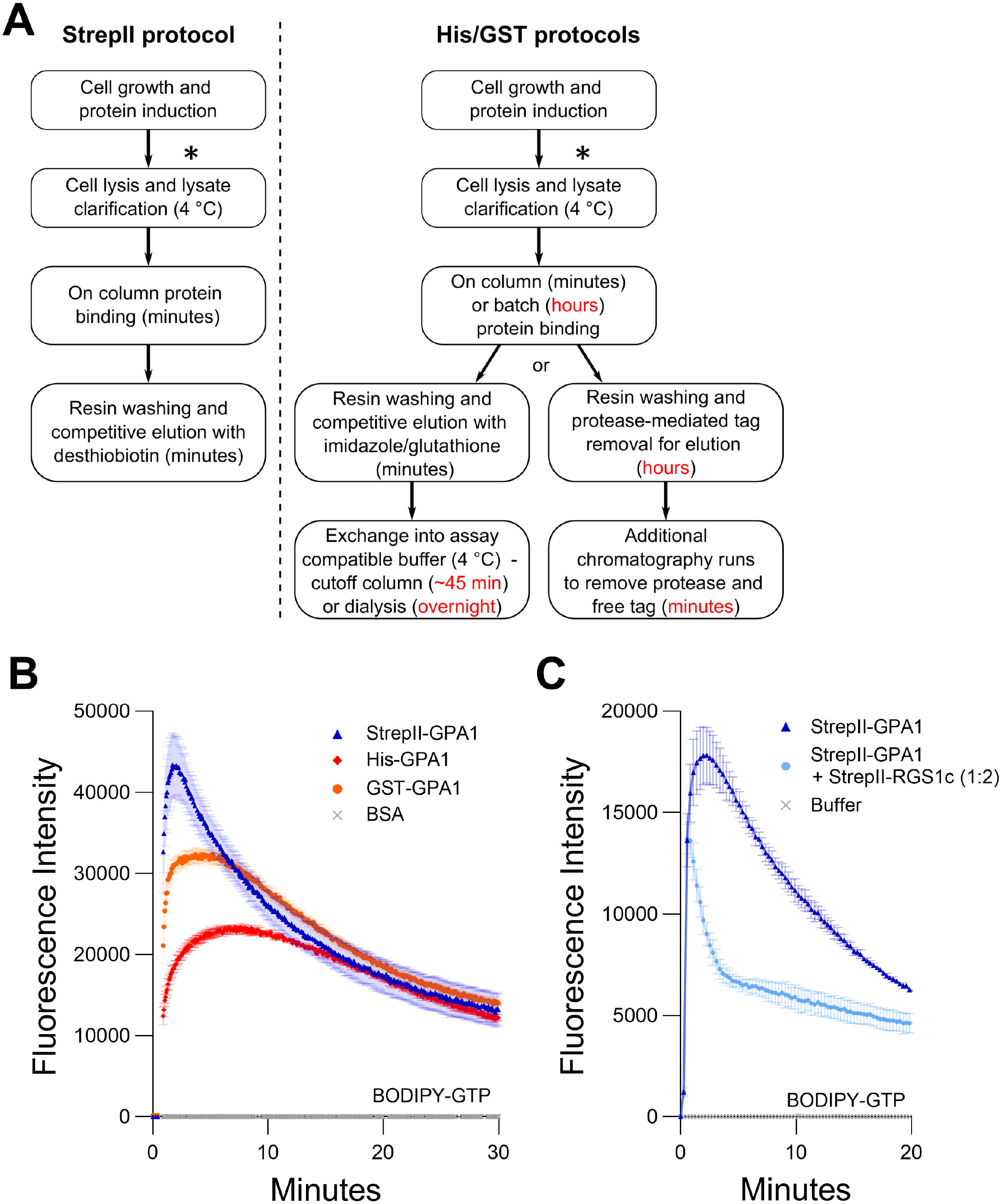
StrepII-tagged protein displays higher enzymatic activity than GPA1 proteins purified using other methods. **A.** Comparison of the purification schemes of StrepII-GPA1 vs. His-/GST-GPA1. Red text indicates points at which timing is extended in traditional protocols. The asterisks (*) indicate a potential stopping point at which cell pellets can be frozen and stockpiled for future purifications. **B.** Comparison of BODIPY-GTP binding and hydrolysis of GPA1 isolated by StrepII-, His- and GST-tag purification procedures. The comparison between StrepII-GPA1 and His-GPA1 was performed five times, while GST-GPA1 was included twice, as the low yield of GST-GPA1, shown in Table S3, precluded extensive characterization. **C.** BODIPY-GTP binding and hydrolysis data for StrepII-GPA1 ±StrepII-RGS1 (cytosolic domain - StrepII-RGS1c). In **B** 100 nM protein was used, with bovine serum albumin (BSA) at equimolar concentration as a negative control. In **C** 90 nM StrepII-GPA1 was used ± 180 nM StrepII-RGS1c. The assay presented in panel **C** was performed six times with similar results; each instance utilized independently purified proteins. To generate the depicted fluorescence intensity values, the raw data values of two (panel **C**) or three (panel **B**) technical replicates for each protein were baseline-corrected using the mean replicate values for the BSA negative control (panel **B**) and averaged, therefore the BSA trace appears near the x-axis. Panel **C** was treated similarly using a buffer control as the negative control. The data are graphically presented as the mean ± SEM and values below zero are not plotted.

### SYPRO Orange assays

We adapted the protein unfolding assay of Biggar et al. [47] to assess protein stability over time at 25 °C. Protein was diluted to 400 nM or 600 nM in “EB Base” supplemented with 5 mM MgCl_2_ and nucleotides as indicated with 5x SYPRO Orange dye (Thermo Fisher Scientific #S6650 - 5000X stock). Forty μl per reaction was aliquoted into wells of a FLUOTRAC 200 96 well half area plate (Greiner Bio-One, 675076), which was loaded into a Synergy Neo2 multimode reader (Biotek). Fluorescence was monitored for the indicated length of time with monochromators set to 470/20 nm excitation and 570/20 nm emission with a gain setting of 100 and a kinetic interval of 5 to 10 seconds. For assays that included a refolding component, Synergy Neo2 injectors were used for the addition of 10 µM GDP (Millipore Sigma, G7127) or 10 µM GTPγS (Millipore Sigma, 10220647001) at the indicated time points.

### Data analysis

BODIPY assays represent the average of 3 technical replicates and were repeated 2-8 independent times (independent biological replicates) with the following exceptions: samples in Figure S1C were assayed in duplicate due to the time constraints of assaying an unstable protein in well-mode and GNAO1^GPA1hel^ was assayed in duplicate in Figure 6B due to yield constraints. SYPRO Orange assays represent the average of 2-3 technical replicates and were repeated 2-6 independent times. Instrument-collected raw data were imported into GraphPad Prism (v10.3) for analysis, baseline corrected as indicated in each figure legend, and graphically presented as the mean ± SEM for all timepoints, except for those in Figures 2C and 2D, where error bars would obscure some trends. Biochemical reaction rates were estimated by hand-adjusting parameters for one-phase association (Y=Y0+(Plateau-Y0)*(1-e^-k*x^)), one-phase decay (Y=(Y0-Plateau)*e^-k*x^+Plateau), and plateau with one-phase decay (Y=IF(x<X0, Y0, Y0+(Plateau-Y0)*(1-e^(-k*(x-X0))^)) models and cross-checking for convergence where appropriate (Prism v10.3). First-order kinetics models were employed following the precedent of Jones et al. [33], Johnston et al. [48], McEwen et al. [49], Eberth et al. [50], Lin et al. [51] and Hewitt et al. [3]. The estimated biochemical rate data and goodness of fit R^2^ values are available in Table S2. Data for Figures 2C and 2D within a span of five seconds before and one second after nucleotide injection were subjected to outlier identification using the robust nonlinear outlier identification module provided in the Prism software, with a 1% false discovery rate. After this analysis, two time points each were removed from the Figure 2C T=20 and T=30 assays, and 2-5 time points each were removed from the T=15, 20, and 30 assays in Figure 2D.

**Figure 2.**
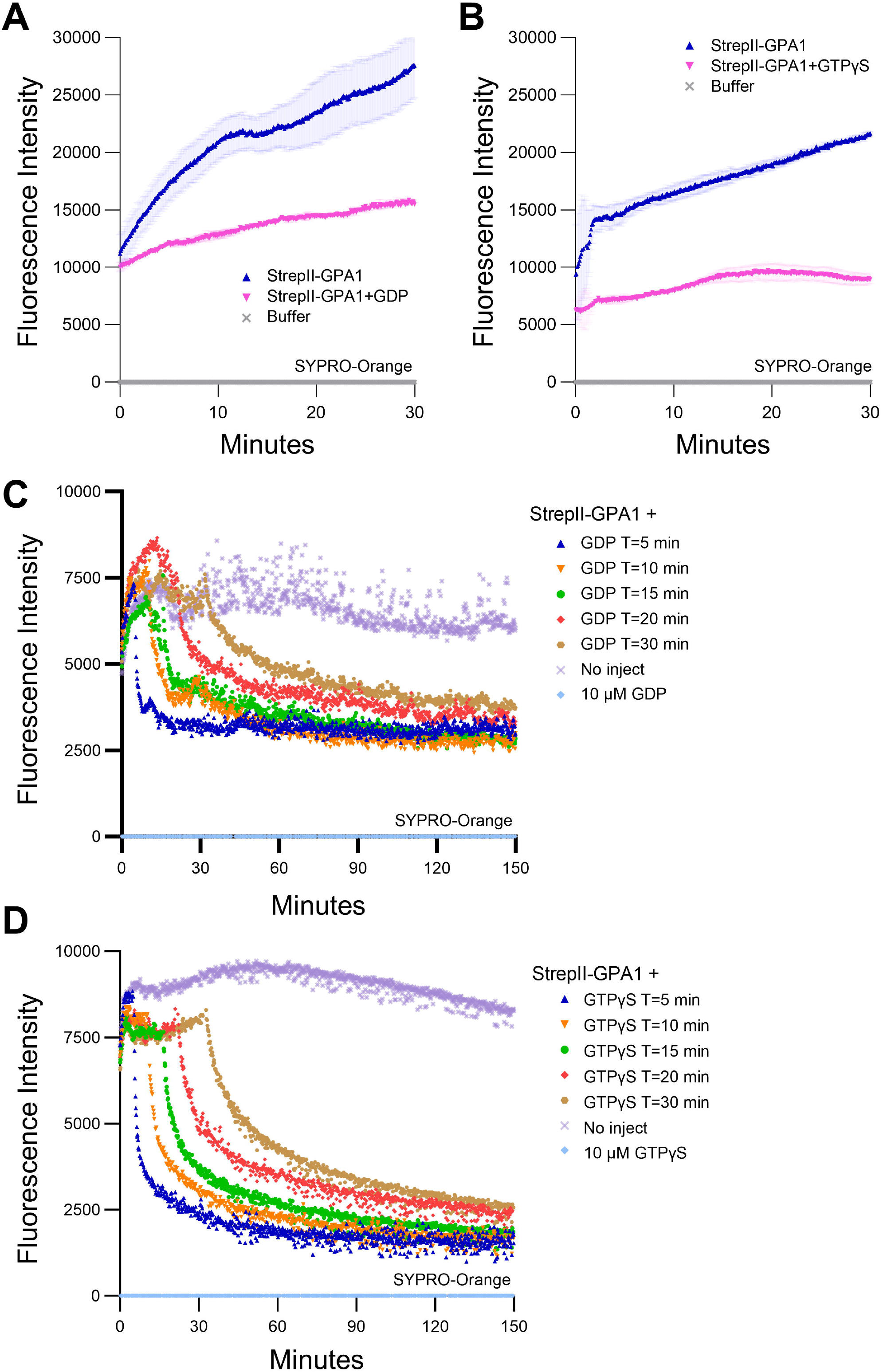
GPA1 displays temperature- and time-dependent loss of activity. **A-B.** SYPRO Orange protein unfolding assays on GPA1 conducted at 25 °C in the absence of additional nucleotides, compared to GPA1 supplemented with **A.** 125 µM GDP or **B.** 125 µM GTPγS. **C-D.** SYPRO Orange protein unfolding/refolding assays on GPA1 conducted at 25 °C with **C.** 10 µM GDP or **D.** 10 µM GTPγS injected at T= 5, 10, 15, 20 or 30 minutes, compared to no injection controls. 600 nM protein was used in SYPRO Orange assays. The assay presented in panel **A** was performed three times with similar results, the assay in panel **B** seven times with similar results and the assays in **C** and **D** six and five times, respectively, with similar results; each instance utilized independently purified proteins. To generate the depicted fluorescence intensity values, the raw data replicate data analyzed were baseline-corrected using the mean replicate values for buffer (panels **A** and **B)**, or 10 µM GDP (panel **C**), or 10 µM GTPγS (panel **D**) controls, which therefore appear near the x-axis. The data are graphically presented as the mean ± SEM in panels **A** and **B**, only the means shown in panels **C** and **D** for clarity, and values below zero are not plotted. All traces in panels **A** and **B** represent the mean of two technical replicates. All traces in panels **C** and **D** represent the mean of three technical replicates, with the exception of the panel **C** T=10 trace, for which one of the three replicates was excluded due to an abrupt artifactual mid-course rise in signal.

**Figure 3.**
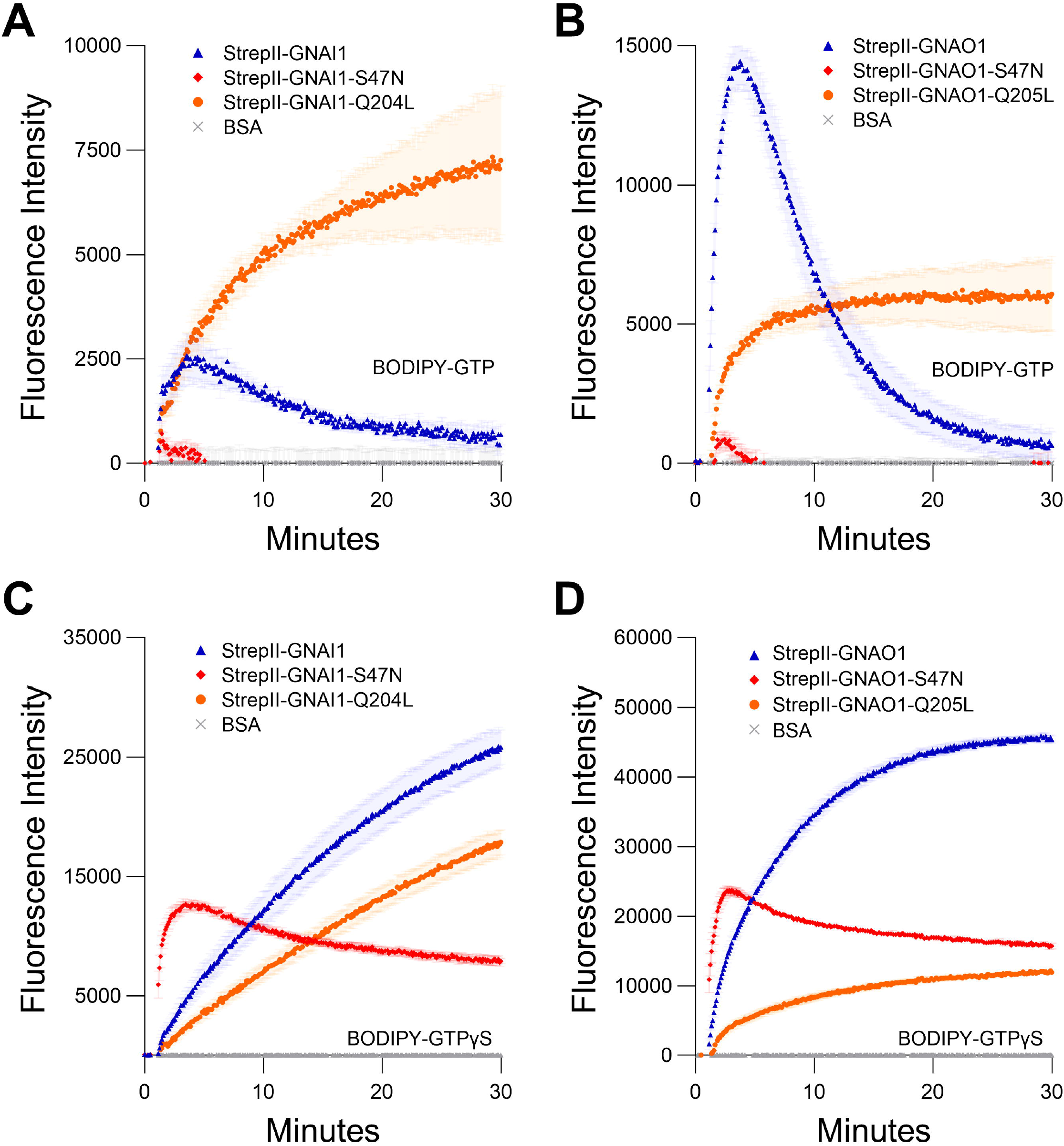
Comparison of StrepII-GNAI1 and StrepII-GNAO1 to their dominant negative (S47N) and constitutively active (Q204L/Q205L) mutants. **A-B.** BODIPY-GTP binding and hydrolysis curves of **A.** GNAI1 or **B.** GNAO1, with corresponding mutants. **C-D.** BODIPY-GTPγS binding curves of **C.** GNAI1 or **D.** GNAO1, with corresponding mutants. Note: for all graphs, wild-type = blue, S47N = red, and Q204L/Q205L = orange. The assays presented in this figure were all performed twice using independently purified proteins with similar results. To generate the depicted fluorescence intensity values, the raw data values of three technical replicates for each protein were baseline-corrected using the mean replicate values for the BSA negative control and averaged, therefore the BSA traces appear near the x-axes. The data are graphically presented as the mean ± SEM and values below zero are not plotted.

**Figure 4.**
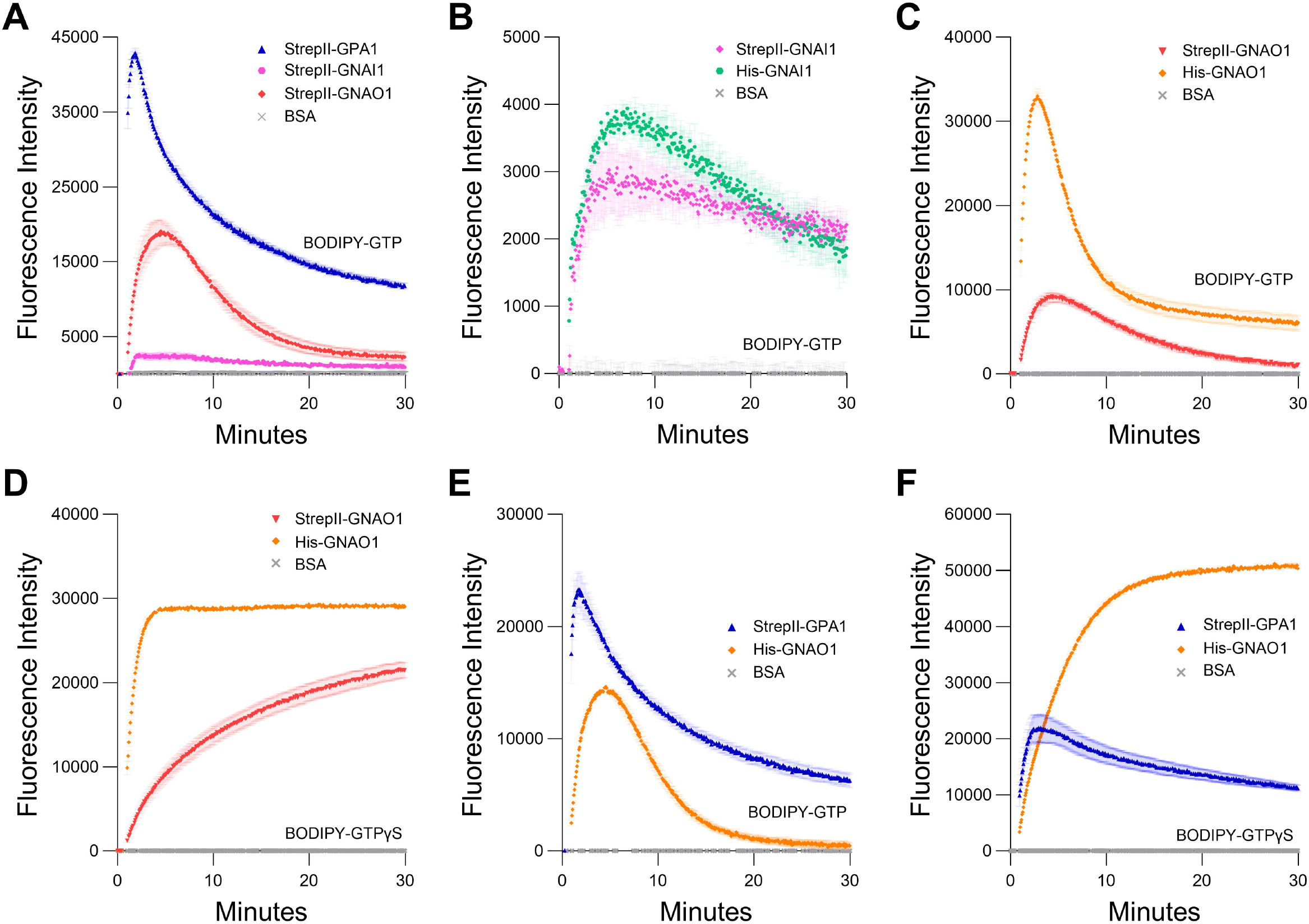
Comparison of GPA1 activity to GNAI1 and GNAO1 activity. **A.** BODIPY-GTP binding and hydrolysis curves of StrepII-GPA1, StrepII-GNAI1 and StrepII-GNAO1. **B-C.** BODIPY-GTP binding and hydrolysis curves of **B.** StrepII-GNAI1 vs. His-GNAI1 and **C.** StrepII-GNAO1 vs. His-GNAO1. Note that the low activity of GNAI1, as evident in panel **A**, results in a y-axis scale difference for panel **B** that increases the visibility of noise and minor differences in GNAI1 activity. **D.** BODIPY-GTPγS binding curves of StrepII-GNAO1 vs. His-GNAO1. **E-F.** Comparison of enzyme kinetics of StrepII-GPA1 vs. His-GNAO1. **E.** Binding and hydrolysis of BODIPY-GTP or **F.** binding of BODIPY-GTPγS. Gα proteins were used at 200 nM protein in panel B and 130 nM protein in panels **C** and **D**, while 100 nM protein was used in panels **A**, **E** and **F**. The assays presented in this figure were performed two or three times using independently purified proteins with similar results. To generate the depicted fluorescence intensity values, the raw data values of three technical replicates for each protein were baseline-corrected using the mean replicate values for the BSA negative control and averaged; therefore, the BSA traces appear near the x-axes. The data are graphically presented as the mean ± SEM and values below zero are not plotted.

**Figure 5.**
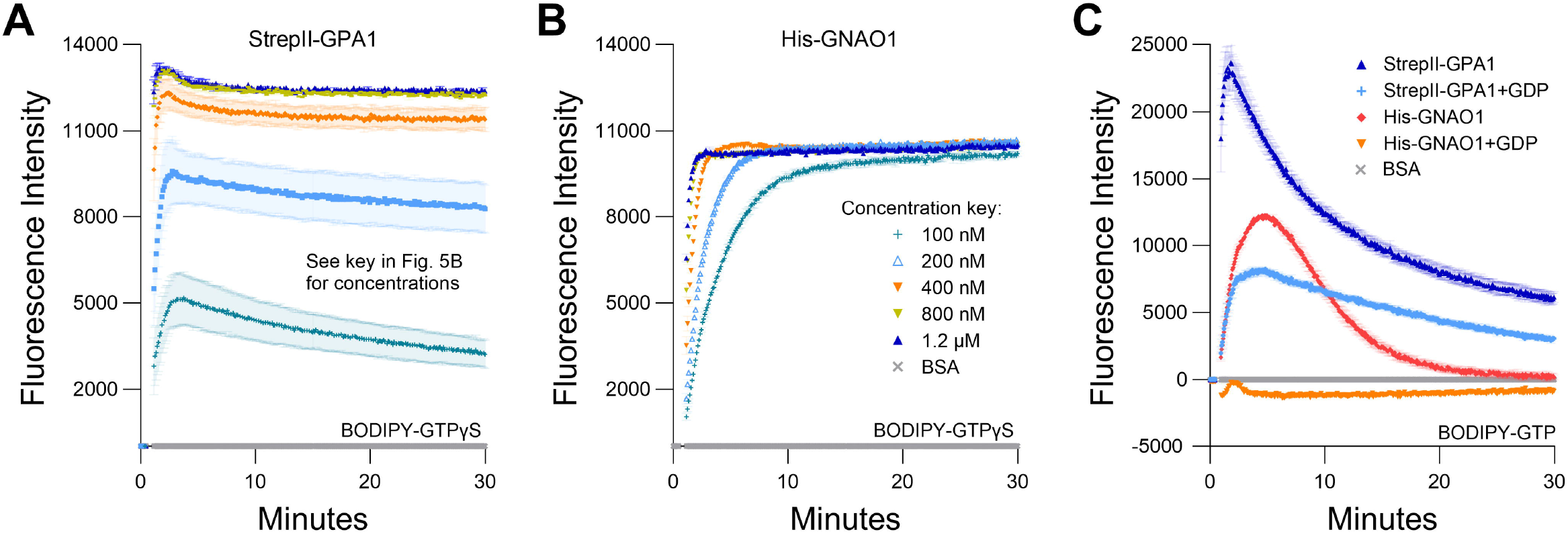
Saturation of BODIPY-GTPγS binding occurs at lower protein concentrations for GNAO1 than GPA1. **A-B.** Concentration-dependent kinetics and maximal binding of 50 nM BODIPY-GTPγS by **A.** StrepII-GPA1 or **B.** His-GNAO1. **C.** Comparison of binding and hydrolysis of BODIPY-GTP by StrepII-GPA1 vs. His-GNAO1 ±10 µM GDP. Gα proteins were used at 100 nM protein in panel **C**. The assays presented in this figure were performed two or three times using independently purified proteins with similar results. To generate the depicted fluorescence intensity values, the raw data values of two (panels **A** and **B**) or three (panel C) technical replicates for each protein were baseline-corrected using the mean replicate values for the BSA negative control and averaged, therefore the BSA traces appear near y=0. The data are graphically presented as the mean ± SEM and values below zero are not plotted, except in panel **C** to show the presence of the His-GNAO1+GDP samples.

**Figure 6.**
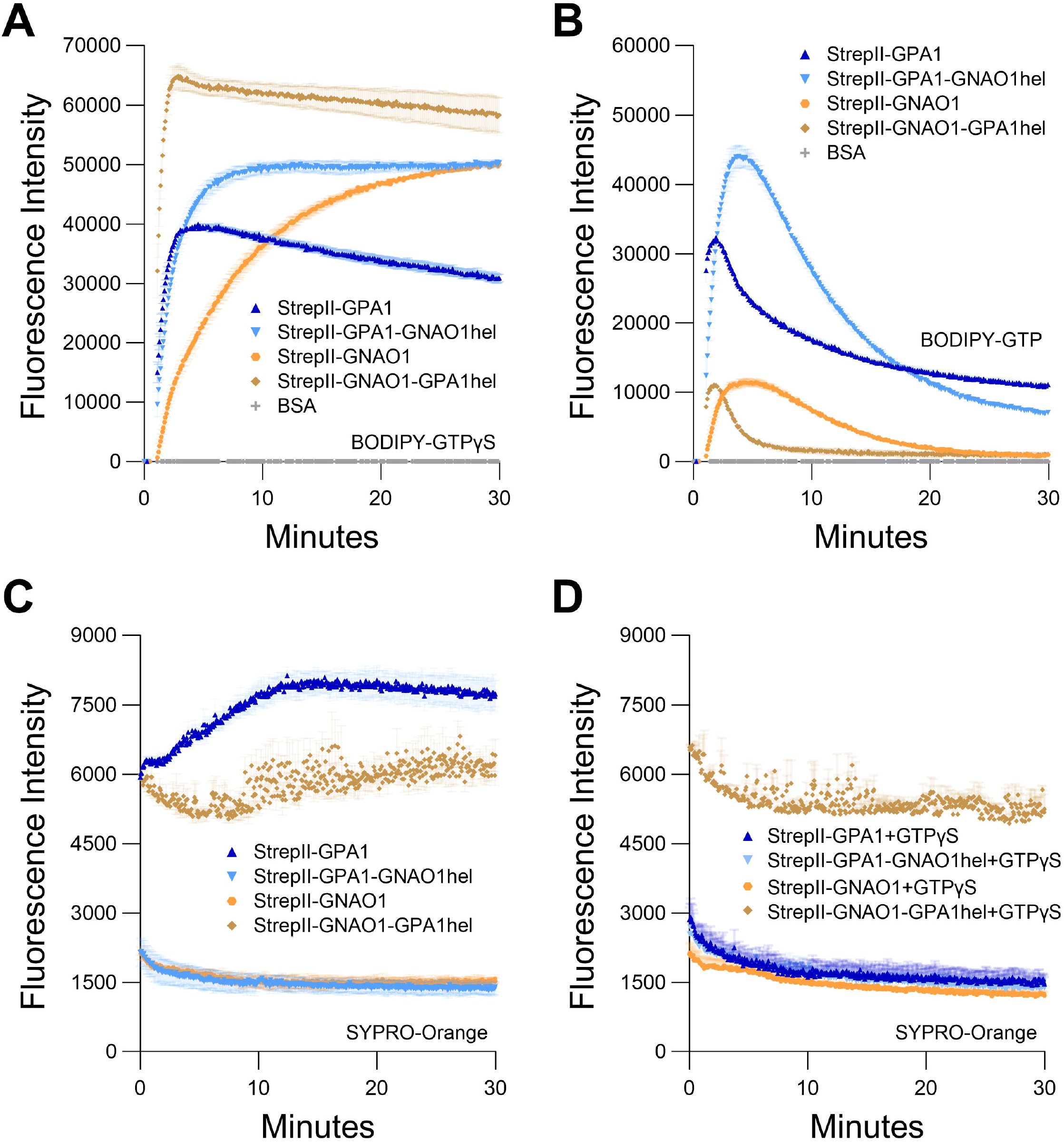
Helical domain swap between GPA1 and GNAO1. Regions encoding the helical domains of GPA1 (residues 68-188) and GNAO1 (residues 63-177) were reciprocally swapped by overlap-extension PCR and the resultant constructs were all expressed with dual StrepII-tags to eliminate the tag as a variable. **A-B.** Curves of GPA1, GNAO1 and helical domain swaps for **A.** BODIPY-GTPγS binding and **B.** BODIPY-GTP binding and hydrolysis. **C-D.** SYPRO Orange protein unfolding assays conducted at 25 °C with GPA1, GNAO1, GPA1^GNAO1hel^ or GNAO1^GPA1hel^ **C.** in the absence of supplementation with additional nucleotides, or **D.** in the presence of 10 µM GTPγS. 100 nM protein was used in BODIPY assays and 400 nM protein in SYPRO Orange assays. The assays presented in this figure were performed two or three times using independently purified proteins with similar results. To generate the depicted fluorescence intensity values, the raw data values of two or three technical replicates for each protein were baseline-corrected using the mean replicate values for the BSA negative control and averaged, therefore the BSA traces appear near the x-axes in panels **A** and **B,** and values below zero are not plotted. The data for panels **C** and **D** are directly presented as the replicate means without error bars for clarity.

## Results

### Rapid purification of recombinant GPA1 via tandem StrepII tags yields higher activity *in vitro*

The dual StrepII tag consists of tandem StrepII tags separated by a flexible linker. The original Strep tag was identified as a streptavidin binding tag that could be used to isolate recombinantly expressed antibodies [52]. Both the Strep tag as well as streptavidin were further engineered to form a StrepII-Streptactin system with increased affinity, and the option of either N-terminal or C-terminal tagging [53,54]. As the resin-conjugated Streptactin used to purify StrepII-tagged proteins exists in a tetrameric state, the use of two tandem StrepII tags separated by a linker was subsequently employed to further increase binding affinity via the avidity effect while still allowing efficient competitive elution [53] in a buffer compatible with many downstream *in vitro* assays. These characteristics offer the advantages of rapid purification of highly pure protein without the need for post-elution processing (Fig. 1A). We therefore utilized dual N-terminal StrepII tags separated by a linker sequence (SGGSGTSGGSA), similar to the linker used in the Twin-Strep-tag (GGGSGGGSGGSA) [53], to purify GPA1, the sole canonical Gα subunit from Arabidopsis. We adapted the base pGEX vector backbone to include N-terminal dual-StrepII tags, thrombin and TEV protease sites, a multiple cloning site, and the option of a C-terminal FLAG tag in a new vector we named pSTTa. GPA1 expressed from pSTTa was purified and observed to be highly pure (Fig. S1A). After numerous trials of multiple growth and induction protocols, we found that use of BL21 (DE3) cells cultured in high salt LB (HSLB) media resulted in the highest and most consistent yields of StrepII-GPA1, so we utilized HSLB for growth and induction in all subsequent experiments.

We first compared StrepII-GPA1 purification to existing His-GPA1 and GST-GPA1 methods (Fig. 1A). To circumvent detrimental overnight dialysis steps that risk protein aggregation [55], we prepared His-GPA1 and GST-GPA1 fusion proteins fresh and as rapidly as possible, utilizing 10 kDa molecular weight cut-off centrifugal filter units for post-elution buffer exchange into “EB Base”, the buffer used for StrepII-GPA1 elution but lacking desthiobiotin. When purified side-by-side with StrepII-GPA1, the buffer exchange steps applied to the His-GPA1 and GST-GPA1 proteins added approximately 45 minutes of additional handling (Fig. 1A), which was performed on ice and in a 4 °C refrigerated centrifuge, while the StrepII-GPA1 sample was kept on ice. When separated by SDS-PAGE, the StrepII-GPA1 protein was obviously more pure than the His-GPA1 or GST-GPA1 preparations under these rapid purification conditions (Fig. S1B). After multiple rounds of side-by-side purification it was notable that the yield of the StrepII-GPA1 protein was consistently ∼2-fold and ∼8-fold higher than that of His-GPA1 or GST-GPA1, respectively (Table S3).

In a classic in vitro assay, fluorescent signal increases as Gα proteins bind BODIPY-conjugated GTP and decreases when GTP hydrolysis outpaces GTP binding, due to resultant partial fluorophore re-quenching [46,49,56–59]. We found that freshly purified StrepII-GPA1 exhibits a characteristic BODIPY-GTP kinetic curve indicative of rapid GTP binding and rapid GTP hydrolysis (Fig. 1B). Despite comparable affinity purification protocols and rapid handling at cold temperatures for the post-purification buffer exchange steps, the His-GPA1 and GST-GPA1 proteins displayed lower apparent binding (39% and 77% of the StrepII-GPA1 rate of 3.139 min^-1^) and hydrolysis (15% and 61% of the StrepII-GPA1 rate of 0.094 min^-1^) activities than the StrepII-GPA1 protein (Fig. 1B, Table S2). As peak fluorescence of BODIPY-GTP is a function of net binding and hydrolysis, another interpretation for the lower activity observed for His- and GST-fusions in Figure 1B is that GTP binding rate is unaltered between the proteins, but GTP hydrolysis is faster in the His- and GST-fusions. We therefore assayed binding to the non-hydrolyzable BODIPY-GTPγS using rapid sampling in well mode for 30 seconds to assess the initial relative GTPγS-binding rates. Indeed the StrepII-GPA1 protein displayed appreciably faster BODIPY-GTPγS binding compared to the His-GPA and GST-GPA1 preparations (Fig. S1C), consistent with the interpretation that the StrepII-GPA1 preparation displays higher activity.

An alternative explanation for the increased apparent activity of the StrepII-GPA1 protein compared to His-GPA1 and GST-GPA1 (Fig. 1B) is that a co-purifying contaminant in the StrepII-GPA1 preparations displays GTP-binding and hydrolysis activities. Though the StrepII purifications commonly yield highly pure protein, we almost always observed a minor contaminating band slightly larger than 70 kDa in our GPA1 elutions (Fig. S1A). To verify the lack of GTP binding by any co-purifying contaminant, we expressed and purified a GPA1^S52N^ mutant that, as observed previously for a GPA1^S52C^ mutant [46], does not bind GTP. StrepII-GPA1^S52N^ displayed no BODIPY-GTP or BODIPY-GTPγS-binding activity (Fig. S1D and E), confirming that the binding and hydrolysis observed for StrepII-GPA1 was solely a result of GPA1 activity. Mass spectrometric identification was performed on the ∼70 kDa protein and it was found to correspond to *E. coli* DnaK, which is not expected to display GTP binding activity [60].

We further functionally validated our StrepII-GPA1 protein using RGS1, which is a key biochemical regulator of GPA1. Arabidopsis RGS1 encodes a protein with seven transmembrane spanning domains at the N-terminus and a cytosolic RGS domain at the C-terminus that stimulates the GTPase activity of GPA1. Previous studies of RGS1 activity have therefore utilized only the cytosolic RGS domain to study its effect on GPA1 *in vitro* [61,62]. We recombinantly produced the RGS1 cytosolic domain utilizing the pSTTa vector and tested its effect on StrepII-GPA1 to confirm: i) that RGS1 is also amenable to rapid purification via dual StrepII-tags, and more importantly; ii) that the StrepII-tagging approach for GPA1 did not disrupt GPA1 interaction with the primary regulator, RGS1. The addition of StrepII-RGS1 to StrepII-GPA1 in a BODIPY-GTP assay strongly promoted the hydrolysis of GTP by GPA1 (Fig. 1C), indicating both i and ii are true.

We note that when our assays were run in well mode, the variation between technical replicates of the StrepII-GPA1 samples was high with a drop in signal observed from wells measured later within the same run (Fig. S1F), as reflected by the large error bars in Figure S1C and S1E. To establish if this variability was intrinsic to StrepII-GPA1, or an artifact of the delay between measurement of samples in a well mode assay, we performed a well mode assay in which three StrepII-GPA1 technical replicates were measured sequentially, prior to the negative control reactions (Fig. S1G). This rapid sampling resulted in greatly reduced replicate-to-replicate variability as displayed by the smaller error bars in Figure S1G compared to Figure S1C. These results also were consistent with a time-dependent loss of activity for GPA1, a phenomenon we investigated further through assessment of GPA1 stability.

### GPA1 stability

To investigate the underlying cause of *in vitro* GPA1 activity loss observed during a sampling timecourse (Fig. S1F), we assessed StrepII-GPA1 conformational stability utilizing SYPRO Orange fluorescence. SYPRO Orange fluorescence increases upon interaction with hydrophobic regions of a protein, which will have greater accessibility upon protein unfolding. We observed that at our standard BODIPY-GTP binding assay temperature of 25 °C, GPA1 protein displayed a steady increase in SYPRO Orange fluorescence over the course of 30 minutes, indicative of protein unfolding (Fig. 2, A and B) as the cause of the activity loss. This increase in protein instability was largely counteracted by incubating GPA1 with excess unlabeled GDP (Fig. 2A) or GTPγS (Fig. 2B). Indeed, many Gα purification protocols include excess concentrations of GDP in buffers to stabilize the protein and prevent aggregation [55] during protracted purification protocols such as those that utilize proteolytic cleavage or buffer exchange by techniques including overnight dialysis ([3,33,34,40,46,63–69] and Fig. 1A). We predicted that, given the greatly reduced processing time, our streamlined StrepII purification protocol would be suitable for purification of active Gα proteins without the need for supplementation with excess GDP. Gα proteins purified from cell lysates are expected to remain at least partially GDP loaded in the short term from production in *E. coli*, however, over time if not provided with excess nucleotide, GDP dissociation will result in protein destabilization. Indeed when assayed fresh our StrepII-GPA1 preparations displayed higher peak signals and increased BODIPY-GTPγS (Fig. S2A, Tables S2 and S4) and BODIPY-GTP (Fig. S2B, Tables S2 and S4) binding activities compared to preparations that had been freshly eluted in the presence of 10 µM GDP. These data confirmed that when prepared with our rapid purification protocol and assayed immediately post-quantification, StrepII-GPA1 could be assayed without GDP supplementation.

To correlate GPA1 unfolding with loss of activity over time in the absence of excess nucleotides, we assayed BODIPY-GTP binding with or without a 30-minute 25 °C preincubation, analogous to the SYPRO-Orange assay timeline in Figure 2A. After the 30-minute 25 °C preincubation GPA1 displayed clearly reduced BODIPY-GTP binding activity (Fig. S2C), with an association rate (2.123 min^-1^) only 57% that of freshly eluted StrepII-GPA1 (3.706 min^-1^) (Table S2). The inclusion of excess (10 µM) GDP, with or without 30-minute preincubation at 25 °C, reduced maximum net binding signals by ∼75% (Fig. S2C, Table S4). Therefore, the presence of excess (10 μM) GDP prevented temperature-dependent loss of activity (Fig. 2A and B); however, competition assays revealed that GDP also suppressed BODIPY-GTP binding (Fig. S2A-C), which could lead to underestimates of binding and hydrolysis rates.

We then investigated the extent to which nucleotides could stimulate GPA1 refolding. SYPRO Orange assays at 25 °C were performed as before, except GDP or GTPγS were injected at different time points during the first 30 minutes. Notably, with multiple samples assayed in triplicate the setup time of the assay took several minutes, and by the time measurements were initiated it is presumed some portion of the early unfolding phase displayed in Figures 2A and 2B had occurred. Addition of GDP (Fig. 2C) or GTPγS (Fig. 2D) during the course of the assay stimulated a partial reduction in SYPRO Orange signal, indicative of protein refolding induced by the nucleotide. The reduction in SYPRO Orange signal was generally more rapid and occurred at a greater magnitude when the nucleotides were injected at earlier time points, e.g. T=5 minutes vs. T=30 minutes (Table S2). These results indicate that GPA1 unfolding is partially reversible by nucleotide addition, but the extent and speed to which GPA1 can refold is time-dependent.

We next examined if StrepII-GPA1 was amenable to storage overnight at 4 °C. Fresh StrepII-GPA1 (Fig. S2D) displayed 19% greater maximal GTP binding, 46% faster binding kinetics, and a 66% faster hydrolysis rate compared to StrepII-GPA1 after overnight storage at 4 °C (Tables S2 and S4), indicating that overnight storage causes an overall loss of activity. Given the stability issues of GPA1 outlined above, we hypothesized that storage at −80 °C and freeze-thawing also has an impact on activity of GPA1 in the absence of GDP supplementation. Therefore, we purified StrepII-GPA1, aliquoted the preparation into various concentrations of either glycerol or sucrose as a cryoprotectant [70], snap froze these fractions with liquid N_2_ and stored the proteins for three weeks at −80 °C. Upon thawing and assaying the protein, we observed that GPA1 frozen with sucrose as a cryoprotectant displayed higher peak BODIPY-GTP binding activity than GPA1 frozen with glycerol (Fig. S3A). We then repeated the assay to compare the binding and hydrolysis curves of StrepII-GPA1 protein stored with the optimal sucrose concentration, 8.33%, vs. the optimal glycerol concentration, 10%. Figure S3B shows StrepII-GPA1 frozen with sucrose exhibited 30% more rapid BODIPY-GTP binding and 123% faster hydrolysis compared to StrepII-GPA1 stored with glycerol (Table S2). StrepII-GPA1 prepared either fresh or frozen with 8.33% sucrose displayed similar BODIPY-GTP binding and hydrolysis curve shapes (Figs. 1B vs. S3B). This difference suggests the use of glycerol as a cryoprotectant could result in an underestimate of the peak net hydrolysis rate of GPA1 preparations.

With the exception of the frozen storage experiments reported above, all of the data presented herein are results from freshly isolated protein. In the following comparative experiments, freshly isolated GPA1, GNAI1, and GNAO1 proteins were stored on ice in a 4 °C refrigerator during post-elution quantification steps, the reaction mixes were prepared on ice, and once aliquoted to plates were subjected to no more than 2 minutes of temperature equilibration immediately prior to initiating assays, to mitigate loss of activity.

### Comparison of GPA1 to human GNAI1/GNAO1

Arabidopsis GPA1 bears high structural similarity to human GNAI1, which has provided a rationale for previous biochemical comparisons of GPA1 with GNAI1 such as those conducted by Jones et al. [33]. We therefore sought to extend our newly optimized recombinant Gα purification protocol to GNAI1. We cloned GNAI1 into pSTTa using both codon harmonized (GNAI1ch) and wild-type (GNAI1wt) sequence (Fig. S4, A and B). We found that proteins derived from the two constructs were essentially interchangeable as side-by-side comparisons showed the GNAI1wt and GNAI1ch proteins did not differ in yield or purity (Fig. S4C), in BODIPY-GTP binding and hydrolysis (Fig. S4D), or in BODIPY-GTPγS binding (Fig. S4E). To expand upon the previous in-depth comparison of GPA1 with GNAI1 [33], we prepared a construct for another human Gα_i_ subfamily member, GNAO1. GNAO1 has been shown to display considerably faster kinetics than GNAI1 [58] and in a brief comparison to GPA1, displayed more similar kinetic properties to GPA1 [71] than GNAI1 did. On a sequence level, the GNAI1 protein shares 38.2% identity and 56.8% similarity with GPA1, while GNAO1 displays 37.0% identity and 54.1% similarity with GPA1. Therefore, it is reasonable to compare GPA1 to both of these mammalian Gα_i_ proteins.

We purified StrepII-GNAI1 and StrepII-GNAO1, and compared their wild-type activities to those of their constitutively active mutants (StrepII-GNAI1^Q204L^/GNAO1^Q205L^ [72]), and to activities of mutants (StrepII-GNAI1^S47N^/StrepII-GNAO1^S47N^) corresponding to the plant nucleotide-free (Figs S1, D and E) GPA1^S52N^ mutant. Both wild-type GNAI1 and GNAO1 proteins displayed GTP-binding and hydrolysis activities, as expected based on previous studies [58]. The net GTP binding activity of GNAI1, reflected by the amplitude of peak fluorescence, was considerably lower (5.6-fold) than that of GNAO1 (Fig. 3, A and B, Table S4). These results demonstrate the StrepII purification protocol is applicable to human Gα subunits. The constitutively active mutants (Q204L/Q205L) displayed slower binding than the wild-type proteins and no hydrolysis activity, as expected, while the S47N mutants displayed no BODIPY-GTP binding activity (Fig. 3, A and B). Surprisingly, the S47 mutants both displayed BODIPY-GTPγS binding activity with initial rates that were faster than was observed for the wild-type GNAI1 and GNAO1 proteins (Fig. 3, C and D). The binding activity, however, showed a peak at 3-4 minutes, followed by a steady decline in signal. Given that the BODIPY fluorophore is covalently attached differently in BODIPY-GTP (ribose ring) and BODIPY-GTPγS (γ-phosphate), inconsistencies in binding results between the two BODIPY reagents could arise from a combination of steric differences of the binding pocket between S47N mutants and the respective locations of the BODIPY fluorophore.

The steady decline of BODIPY-GTPγS binding by the GNAI1 and GNAO1 S47N mutants is unlikely to be due to hydrolysis as GTPγS is considered non-hydrolysable, and no evidence of BODIPY-GTPγS hydrolysis was evident in any of our assays with wild-type GNAI1 or GNAO1. These results are in contrast to the analogous GPA1^S52N^ mutant, which displayed no BODIPY-GTPγS binding (Fig. S1E). We hypothesized that the slow decline in BODIPY-GTPγS fluorescence arose from decreased Mg^2+^ binding. The S47 residue within the G1 motif is important for Mg^2+^ cofactor coordination [73] and since other metal ions are known to inhibit Gα nucleotide binding [74], we routinely utilized trace-metal-free (TMF) grade components to standardize our assays, which explicitly ruled out any effect of extraneously-present divalent ions, including Mg^2+^. Notably, we also show that TMF components are not necessary for basic assays, and our methodology can be performed using standard grade reagents (Fig. S4F). To examine if the S47N mutants do retain some residual Mg^2+^ binding, we performed BODIPY-GTPγS binding assays ±Mg^2+^. In our ±Mg^2+^ assay, both wild-type GNAI1 and GNAO1 and S47N mutants of GNAI1 and GNAO1 displayed a clear requirement for Mg^2+^, with a very low level of BODIPY-GTPγS binding activity observed in the absence of Mg^2+^ (Fig. S4, G and H, Table S4). These data suggest GNAI1^S47N^ and GNAO1^S47N^ do retain some affinity for Mg^2+^ and a requirement for this cofactor in nucleotide coordination. The slow decline in the BODIPY-GTPγS binding may reflect weaker coordination of Mg^2+^ by the mutant proteins.

Next we compared the binding activities of StrepII-GPA1 to StrepII-GNAI1 and StrepII-GNAO1. When compared to StrepII-GPA1, StrepII-GNAI1 displayed a much lower apparent BODIPY-GTP binding peak (6% of StrepII-GPA1), while StrepII-GNAO1 displayed an intermediate activity (44% of StrepII-GPA1) (Fig. 4A, Table S4). As peak fluorescence reflects the net activity of GTP binding and hydrolysis, these initial results were consistent with GPA1 displaying a faster binding and slower hydrolysis rate than GNAI1. GNAO1 displayed greater net BODIPY-GTP binding compared to GNAI1, and an activity more similar to that of GPA1. As the StrepII purification protocol was superior to His purification for GPA1, we characterized StrepII-tagged GNAI1 and GNAO1 in comparison to the commonly used His-tagged GNAI1 and His-tagged GNAO1. We observed that different Gα subunits, even within the Gα_i_ family, display their own optimal tag and purification method. Minor differences in activity were observed between His-GNAI1 and StrepII-GNAI1 (Fig. 4B), indicating that StrepII or His purification is suitable for GNAI1. By contrast, His-GNAO1 displayed higher net BODIPY-GTP binding (1.8-fold) and hydrolysis activities (2.9-fold) than StrepII-GNAO1 (Fig. 4C, Table S2), and the difference was similarly evident for binding of BODIPY-GTPγS (Fig. 4D), indicating the His purification protocol is the more suitable method to assay GNAO1 activity.

Given the above results, we performed side-by-side purifications of StrepII-GPA1 and His-GNAO1, which demonstrated that GPA1 does indeed display a faster GTP binding rate than GNAO1 (3.25 vs 0.80 min^-1^), but the net hydrolysis rates appear not to be as different between plant and human Gα subunits as previously thought [33,48,71], with His-GNAO1 displaying just a 1.8-fold higher hydrolysis rate than StrepII-GPA1 (Fig. 4E, Table S2). To isolate the observed binding rate of the Gα proteins, we performed assays with the non-hydrolyzable GTP analog BODIPY-GTPγS. Indeed StrepII-GPA1 displayed a 7-fold higher initial rate of BODIPY-GTPγS binding than His-GNAO1 (Fig. 4F, Table S2). Contrastingly, the StrepII-GPA1 maximal BODIPY-GTPγS binding signal was 2.3-fold lower than that of His-GNAO1, and rather than plateauing as with GNAO1, the GPA1 BODIPY-GTPγS signal peaked followed by a decrease over time (Fig. 4F, Table S4). Possible reasons for the difference in signal maxima include: i) steric differences of the Gα binding pockets resulting in differential levels of BODIPY fluorophore unquenching upon protein binding, and; ii) inherent instability of GPA1 resulting in a lower apparent binding activity *in vitro*. We believe hypothesis i) is unlikely as the empirically derived crystal structures of GPA1 and GNAO1 are highly similar (Fig. S5A), just as are the structures of GPA1 and GNAI1 (Fig. S5B). We therefore sought to assess the amount of each enzyme necessary to observe saturated and stable binding of 50 nM BODIPY-GTPγS, as a reflection of Gα activity retained *in vitro*. Despite being in excess, 100 nM, 200 nM and 400 nM concentrations of StrepII-GPA1 were unable to attain a maximal binding signal with 50 nM BODIPY-GTPγS. Only 800 nM or 1.2 µM GPA1 displayed a stable binding plateau at the maximal level, with association binding rates of 3.364 and 3.531 min^-1^, respectively (Fig. 5A, Table S2). In comparison, all concentrations of GNAO1 either attained a maximal plateau, or neared maximal fluorescence in the case of 100 nM GNAO1, within the course of our assay (Fig. 5B, Table S4). The necessity for higher GPA1 concentrations in reaching binding saturation reflects the lower stability of GPA1 *in vitro* (Fig. 2, A and B) compared to GNAO1 and GNAI1 (Fig. S5C), but also provides insight regarding the GNAO1>GPA1 signal maxima in Figure 4F. Notably, the maximal levels of BODIPY-GTPγS fluorescence were similar between high concentrations of GPA1 and GNAO1 when assayed side-by-side (Fig. 5, A and B), thereby indicating that steric differences in the binding pockets do not result in markedly different levels of BODIPY fluorophore unquenching and thus refuting hypothesis i) above. Next we compared the ability of excess (10 µM) GDP to suppress binding of 50 nM BODIPY-GTP to 100 nM Gα protein. Peak BODIPY-GTP binding was suppressed 2.9-fold by 10 µM GDP for GPA1; by contrast, BODIPY-GTP binding was almost completely abolished for GNAO1 by 10 µM GDP (Fig. 5C, Table S4). The striking difference in GDP suppression of GTP binding likely reflects a higher relative affinity for GTP than GDP of GPA1, as previously reported [71], and a significantly faster nucleotide exchange rate of GPA1 than GNAO1.

### GPA1-GNAO1 helical domain swaps

It has been previously demonstrated that the helical domain of GPA1 displays a marked level of intrinsic disorder and increased dynamic motion compared to that of GNAI1 [33]. Jones et al. [33] confirmed that a helical domain swap between GPA1 and GNAI1 largely swapped the relative kinetics between the two Gα proteins. In those studies, the GPA1 helical domain conferred rapid spontaneous activation to GNAI1 while the GNAI1 helical domain conferred slower activation to GPA1. Given the more similar activities of GPA1 and GNAO1 (cf. Fig. 4E), we sought to expand upon these analyses to assess if a helical domain swap between GPA1 and GNAO1 would: i) display as strong a difference as the GPA1-GNAI1 domain swap, and ii) confirm that the helical domain of GPA1 is responsible for the poor stability of GPA1. Therefore, we created reciprocal domain swap constructs GPA1^GNAO1hel^ (GPA1 Ras domain fused to the GNAO1 helical domain) and GNAO1^GPA1hel^ (GNAO1 Ras domain fused to the GPA1 helical domain). To not confound any tag/purification effects with the domain swap effects, we utilized our StrepII tagging and purification methods for all proteins. A comparison of BODIPY-GTPγS binding demonstrated that binding rates increased in the following order: GNAO1<GPA1^GNAO1hel^<GPA1<GNAO1^GPA1hel^ (Fig. 6A, Table S2). Beyond this initial binding rate, GPA1 displayed the lowest signal amplitude corresponding to peak binding, while GNAO1^GPA1hel^ displayed the highest signal plateau (Fig. 6A, Table S4). In BODIPY-GTP assays, which integrate GTP binding and hydrolysis, a similar initial trend was largely displayed during the binding phase: GNAO1<GPA1^GNAO1hel^<GNAO1^GPA1hel^<GPA1 (Fig. 6B, Table S2). Once BODIPY-GTP hydrolysis exceeded the binding rate, we observed the following order of maximal net hydrolysis rates: GNAO1=GPA1^GNAO1hel^<GPA1<GNAO1^GPA1hel^ (Fig. 6B, Table S2). Therefore, although the rate of BODIPY-GTPγS binding by GNAO1^GPA1hel^ was the fastest of the four proteins assayed, its peak BODIPY-GTP fluorescent signal was dampened by a rapid switch to net hydrolysis. We then assessed the relative conformational stability of the GPA1, GNAO1, GPA1^GNAO1hel^ and the GNAO1^GPA1hel^ proteins in a SYPRO Orange assay ±10 µM GTPγS (no BODIPY label). As suspected, the GNAO1 helical domain conferred a similar stability to GPA1 as did excess GTPγS (Fig. 6, C and D). As before, GPA1 samples without nucleotide supplementation displayed increased SYPRO Orange signal indicative of protein unfolding, and GPA1 was the only protein in the domain swap assays to display considerable divergence between the ±GTPγS samples (Fig. 6, C and D). Interestingly, at “T=0” of the SYPRO Orange assay, the fluorescence in the absence of nucleotide supplementation was already much higher for GPA1 than for GNAO1 or GPA1 supplemented with GTPγS (Fig. 6, C and D). We note that all of these samples were prepared on ice in duplicate, pipetted into the assay plate and loaded into the plate reader; a process that took ∼4 minutes for the number of samples in Figures 6C and 6D. To investigate the difference at “T=0” of the assays comparing multiple samples, we performed a 1 vs. 1 assay comparing single wells of GPA1 vs. GPA1 +10 µM GDP. This assay can be initiated in seconds and allowed us to monitor SYPRO Orange fluorescence almost immediately after removal from ice. Indeed, in this rapid assay, the initial fluorescence levels were similar between the samples before a steady rise in fluorescence signal was observed in the GPA1 alone reaction (Fig. S5D). The initial similarity of fluorescence between ±nucleotide samples was quite similar to the results shown in Figures 2A and 2B, which were assays run on an intermediate scale compared to the large assays in Figures 2C, 2D, 6C and 6D, and the small scale assay in Figure S5D. GNAO1^GPA1hel^ in the large scale assay also displayed a higher initial value of SYPRO Orange fluorescence, and noticeably more signal variation between timepoints, though it should be noted that this noise-like variation was not always observed for GNAO1^GPA1hel^ (Fig. S5E). Unlike GPA1, the addition of GTPγS did not repress the T=0 high fluorescence values for GNAO1^GPA1hel^, yet the fluorescence signals for GNAO1^GPA1hel^ did not rise as rapidly through the assay as they did for GPA1 in the absence of GTPγS (Fig. 6, C and D). These traits appear consistent with GNAO1^GPA1hel^ achieving a stable but different conformation than the other Gα proteins assayed in Figure 6; a conformation that is seemingly characterized by increased surface accessibility of hydrophobic residues for SYPRO Orange binding, but with strong BODIPY-GTPγS binding as evident in Figure 6A, Table S2 and Table S4, indicative of a highly active protein. A decrease in BODIPY-GTPγS fluorescence for GNAO1^GPA1hel^ was observed after peak signal had been reached (Fig. 6A), indicating a loss of activity over the course of the assay, similar to the loss observed for GPA1. However, wild-type GPA1 displayed a lower peak of BODIPY-GTPγS fluorescence, indicating a more rapid loss of activity for GPA1 than GNAO1^GPA1hel^, consistent with the more immediate unfolding observed for GPA1 in Figure 6C. In summary, for GPA1 *in vitro*, unfolding at room temperature during plate setup and then at 25 °C upon assay initiation in the plate reader began almost immediately and was evident on a scale of seconds to minutes (Fig. S5D), further underscoring the need to use a rapid purification protocol. Interestingly, our domain swap assays indicate that neither the Ras nor the helical domain alone accounts for the full lack of GPA1 stability, as the magnitude of BODIPY-GTPγS binding indicates the GPA1 helical domain does not destabilize GNAO1 as rapidly as it does GPA1.

## Discussion

Purification of functional recombinant heterotrimeric G protein Gα subunits is integral to understanding their roles in both animals and plants. The former is of importance due to their well-described functions in human health [75], and the latter is important due to G protein involvement in controlling agriculturally important traits [12,76]. We demonstrate for the Arabidopsis Gα subunit, GPA1, that protracted handling and/or storage using the standard protocol of glycerol as a cryoprotectant are detrimental to isolating optimally functional protein (Figs. 1B, 2, S1F, S3A and S3B). Therefore we developed a StrepII-tag expression and purification protocol that allowed rapid on-column binding to isolate highly pure protein for immediate downstream analyses (Fig. 1A). The utilization of an elution buffer compatible with downstream assays, abolishing the need for buffer exchange steps, is a major advantage of the StrepII purification protocol. Even with the rapid StrepII purification protocol, our data were consistent with some loss of GPA1 activity during the purification and assay timeframe, based on the high concentrations of GPA1 required to saturate binding of 50 nM BODIPY-GTPγS, and the BODIPY signal rundown observed at lower concentrations of GPA1 (Fig. 5A, compared to GNAO1 in Fig. 5B). Nonetheless, matched purifications demonstrated that the loss of activity for StrepII-GPA1 was substantially less than that observed for commonly used tags: His-GPA1 or GST-GPA1 (Fig. 1B). It should be noted that inclusion of GDP in Gα protein preparations is well-known to stabilize the protein [55,77], as we also show for GPA1 (Figs. 2A and 2C) and is highly advisable for several applications. Yet, inclusion of excess GDP in elution or storage fractions is not optimal when assaying intrinsic binding affinity as excess concentrations of GDP can compete with GTP for binding to Gα (Figs. 5C and S2A-C). Our rapid purification method was specifically optimized to generate protein that could be assayed prior to substantial loss of activity due to misfolding.

As our method proved to be an improvement over existing protocols for GPA1 purification (Figs 1B and S1C), we applied it to the purification of two closely related human Gα subunits, GNAI1 and GNAO1. We show our method is also applicable to human Gα subunit expression and purification such as for GNAI1 (Fig. 4B). We therefore establish StrepII-mediated purification as an addition to the toolkit of possibilities for recombinant investigation of G proteins. However, His-GNAO1 outperformed StrepII-GNAO1 in our hands (Fig. 4, C and D), reinforcing that tag choice is not universal, and should be optimized for each protein of interest.

We then utilized our newly improved purification protocol to reexamine and further explore the following four questions of interest. 1) Does GPA1 indeed display self-activating properties? 2) Is the balance of GTP-loading of GPA1 further skewed to the active state by slow GTP hydrolysis? 3) What are the functional consequences of mutations to the serine residue important for Mg^2+^ ion coordination in the active site? 4) Given that GNAO1 displays rapid enzyme kinetics compared to GNAI1, but without the loss of stability observed in GPA1, can we employ a domain swap approach between GPA1 and GNAO1 to assess the relative contributions of the Ras vs. helical domains to enzyme function and stability? As GPA1 was sensitive to differences in handling, in all assays we only directly compared proteins prepared fresh side-by-side, and we recommend that as the best practice.

### Re-evaluation of GPA1 enzymatic kinetics

Jones et al. [33] characterized GPA1 as a self-activating Gα protein due to rapid GDP release followed by rapid GTP binding. They also reported slow GTP hydrolysis kinetics. Urano et al. [40] followed this study with confirmation that Gα subunits from evolutionarily distant branches of the plant kingdom also exhibit these properties. However, both studies utilized His-tag purification protocols: Jones et al. purified Gα proteins using a 90-minute batch binding step with post-elution processing steps and compared GPA1 to the slow GTP-binding Gα, GNAI1, and Urano et al. stored purified Gα subunits at −80 °C with glycerol as a cryoprotectant. Though these are standard protocols for Gα purification, with hindsight we now suggest these steps are not optimal for isolation of active GPA1. It should also be noted that both studies included GDP in their elution buffers, which assists in GPA1 stabilization (Fig. 2, A and C) but, depending on concentration, can slow GTP binding and therefore the maximal observable hydrolysis rate (Figs. 5C and S2B-C). As a result we sought to reassess the conclusions drawn from these studies by utilizing our newly developed rapid Gα purification protocol.

All of our data are consistent with the original assertion [33] that GPA1 displays rapid GTP binding (Figs 1B-C, 5C, S1C-G S2 and S4F). Over seven independent BODIPY-GTP experiments, StrepII-GPA1 exhibited a mean association rate of 3.192 ± 0.120 (SEM) min^-1^, which matches the highest protein saturation binding rates for BODIPY-GTPγS (Table S2) and is within the range of previously reported GPA1 association rates (Table S5). Even when compared to GNAO1, which displays much faster binding than GNAI1 (Fig. 4A), GPA1 clearly displays a faster comparative rate of GTP binding (Fig. 4, E and F, Table S2). Furthermore, our analyses of GPA1 stability *in vitro* (Figs 2, 6C-D and S5D), and the inability of moderately excess concentrations of GPA1 to saturate BODIPY-GTPγS-binding (Fig. 5A), suggests that this assessment of GTP binding is still an underestimate due to functional decline under assay conditions. Additionally, the new comparisons to GNAO1 presented in this report reveal the GPA1 hydrolysis rate to be less of an outlier than suggested by previous [33] and current (Fig. 4A) comparisons only to GNAI1. The apparent BODIPY-GTP net hydrolysis rate of GPA1 is only 44% lower than that of GNAO1 (Fig. 4E and Table S2) indicating relatively similar levels of activity. With regard to spontaneous nucleotide exchange, our data in Figure 5C are particularly compelling. In that GDP competition assay, GDP was provided at 10 µM, i.e. 100x in molar excess of the Gα proteins and 200x in molar excess of BODIPY-GTP. This massive overabundance of GDP was sufficient to completely outcompete BODIPY-GTP binding by GNAO1 (Fig. 5C), reflecting the crucial role of GPCR-mediated stimulation in nucleotide exchange for animal Gα subunits [67,78]. Contrastingly, 10 µM GDP was only partially able to suppress GPA1 BODIPY-GTP binding activity (Fig. 5C), consistent with GPA1 displaying a spontaneous nucleotide exchange activity and relatively much higher affinity for GTP than GDP, as previously reported [33,40,71]. Overall, our data indicate that GPA1 does display rapid properties of both nucleotide exchange and GTP binding, but there likely has been underestimation of the GTP hydrolysis activity of GPA1 in the past due to choice of purification protocol. Moreover, side-by-side comparisons with two closely related mammalian Gα proteins, all isolated under optimal conditions (Fig. 4, A and E), reveals that the BODIPY-GTP hydrolysis rate of GPA1 falls within the range of that observed for these animal Gα subunits.

### GNAI1^S47N^ and GNAO1^S47N^ mutants display transient GTP binding

With the advent of affordable mass patient genetic testing, a number of mutations of the equivalent sites to GNAI1^S47^/GNAO1^S47^ and GNAI1^Q204^/GNAO1^Q205^ of multiple Gα subunits have been associated with various medical conditions in ClinVar [79] and Catalogue Of Somatic Mutations In Cancer (COSMIC) [80] databases, as summarized in Tables S6 and S7, respectively. The Q204/Q205 site resides within the G3 motif (one of five G box motifs important for nucleotide binding) of Gα subunits, mutations of which are well-known to impart a constitutively active status upon Gα proteins [72]. Mutations at this site specifically in *GNAQ* and *GNA11* are strongly linked to uveal melanoma [1,81]. The S47 site is relatively less well-understood, though it is a crucial residue within the G1 motif involved in Mg^2+^ cofactor coordination [73]. A Gα_T_ protein, in which a region or subregions of amino acids 215-295 have been replaced with the equivalent GNAI1 residues to facilitate expression and purification, has been utilized and named Gα_T_*. When assaying binding of radiolabeled GTPγS by Gα_T_* chimeric proteins, there was an apparent discrepancy between the results of Natochin et al. [82] who reported S43N and S43C mutants failed to bind GTP, and Ramachandran and Cerione [83] who reported a faster rate of spontaneous GDP-GTPγS exchange for the Gα_T_*^S43N^ mutant compared to Gα_T_*. Our reassessment with real-time BODIPY-GTPγS binding suggests a mechanism by which the discrepancy may be understood. The initial rates of BODIPY-GTPγS binding are faster for GNAI1^S47N^ and GNAO1^S47N^ than for the respective wild-type proteins (Fig. 3, C and D), while retaining Mg^2+^-dependency (Fig. S4, G and H). However, the binding signal of BODIPY-GTPγS by GNAI1^S47N^ and GNAO1^S47N^ decays from an early peak, as shown by the observation that BODIPY-GTPγS signal initially increased, but then gradually decreased 3-4 minutes after binding initiation (Fig. 3, C and D). Thus, the binding signal could be missed and/or washed off if the protein is subjected to protracted handling in a radiolabeled GTPγS binding assay, which may account for the previously reported GTP-binding discrepancy. Our results provide additional insight into the mechanism by which S47 and equivalent position mutations of human Gα subunits may manifest in disease states. Moreover, our results illustrate an advantage of BODIPY assays in facile revelation of real-time kinetics.

### GPA1 instability is conferred by combined effects of the Ras and helical domains, and is not inherently linked to rapid nucleotide binding

As mentioned above, studies from the Jones and Dohlman groups have indicated that the GPA1 helical domain displays both high levels of intrinsic disorder, based on comparisons of the electron density map and atomic displacement parameters of monomers determined by x-ray crystallography, and motion away from the Ras domain, as predicted by molecular dynamics simulations [33,84]. Interdomain motion is a mechanism proposed to potentiate nucleotide exchange [67,78,85,86] and therefore these observations for GPA1 are consistent with its status as a Gα subunit capable of spontaneous nucleotide exchange [33]. As previously established, a domain substitution using the helical domain of GNAI1 conferred slower nucleotide exchange, faster GTP hydrolysis and increased stability to GPA1. Those stability experiments utilized circular dichroism over a temperature gradient of 15-80 °C, and proteins were assayed in the presence of excess GDP. The results of Jones et al. [33] are therefore consistent with a coupling of high activity and low stability through the GPA1 helical domain. To further investigate this phenomenon we utilized a SYPRO Orange fluorescence assay in the presence and absence of additional nucleotides. When incubated at 25 °C without additional nucleotides, GPA1 displayed reduced conformational stability (Figs. 2 and S5D), as expected [55]. We also observed that the enzymatic differences between GPA1 and GNAO1 were less than those between GPA1 and GNAI1 (Fig. 4), though GNAO1 was likely more stable than GPA1 based on the plateau in BODIPY-GTPγS binding signal observed in Figure 5B and similarity of GNAO1 SYPRO Orange fluorescence with and without GTPγS in Figure S5C. We speculated that the helical domain of GNAO1 may confer stability to GPA1 while also allowing the fast GTP binding kinetics of GPA1 to be retained. Indeed this proved to be the case, with GPA1^GNAO1hel^ displaying similarly rapid BODIPY-GTPγS and BODIPY-GTP binding as GPA1 (Fig. 6 A and B, Table S2). When protein stability was assayed, we observed that GPA1^GNAO1hel^ displayed a similar resistance to unfolding at 25 °C as GNAO1, distinguishing it from the less stable GPA1 protein (Fig. 6C). When provided with a molar excess of GTPγS, GPA1 was as stable as GNAO1 and the chimeric Gα subunits (Fig. 6D).

In the reciprocal domain swap, GNAO1^GPA1hel^ displayed rapid BODIPY-GTPγS binding (Fig. 6A) and fast hydrolysis (Fig. 6B). Unexpectedly, GNAO1^GPA1hel^ exhibited a higher basal level of SYPRO Orange interaction than the other Gα proteins, but seemingly with a relatively higher level of protein stability than GPA1 (Fig. 6, C and D). As the GNAO1^GPA1hel^ protein also displays strong enzymatic activity (Fig 6, A and B), we conclude that the GNAO1^GPA1hel^ chimera resides in a conformation characterized by increased SYPRO accessibility, yet still functions as a molecular switch. Therefore, as the GPA1 helical domain alone was not able to confer GPA1-like instability to GNAO1, we propose that interdomain forces contribute to GPA1 instability, and that rapid kinetics and instability can be uncoupled by the use of chimeric domain swaps.

### Future directions

Our results suggest that it will be of interest to further evaluate the GTPase activity of GPA1 in comparison to mammalian Gα subunits. Here we used BODIPY-GTP/-GTPγS to test our new, streamlined purification approach for GPA1, and screen relative G protein activities. We report our purification method as a tool for the community and highlight important contrasts to data from established methods, along with several general consistencies between our data and those of others. We also illustrate an advantage for BODIPY-GTP/GTPγS as it is a real-time method for measurement of direct binding with a sampling rate and processing speed that cannot be matched by traditional radiolabeled nucleotide approaches, as Johnston et al. previously noted [71]. These aspects are particularly useful for proteins with rapid kinetics and low stability *in vitro*. However, we also observed a drawback of the BODIPY labeling approach in the inconsistency observed between BODIPY-GTP and BODIPY-GTPγS binding for GNAI1^S47N^ and GNAO1^S47N^ (Fig. 3, A-D). Conjugation of a fluorophore such as BODIPY to GTP can result in differences in apparent binding compared to unlabeled GTP [87,88], and therefore dictates caution when extending rate estimates to absolute rates. While our study demonstrates greater stability *in vitro* for GNAO1 than GPA1, not all human Gα subunits have been as easy to produce recombinantly as GNAO1. For example, chimeric approaches have previously been required to express Gα proteins in the soluble state, including for mammalians Gα subunits such as GNAT1 [32,35,83], GNA12 and GNA13 [37]. These chimeras integrate short regions of the kinetically particularly slow but easily purified GNAI1 enzyme. Our success with GPA1 purification indicates that our expression and rapid StrepII purification method is worth evaluating for purification of full length recombinant human Gα proteins that are enzymatically active, without the need to resort to chimeric sequence substitutions.

## Acknowledgments

We thank Mr. David Arginteanu for technical assistance and the suggestion to evaluate sucrose as a cryoprotectant. Supported by NIGMS 5R01GM126079 and NIGMS 1R35GM153492 to SMA. The content is solely the responsibility of the authors and does not necessarily represent the official views of the National Institutes of Health.

## Author contributions

TEG and DC: were responsible for conceptualization, performed experiments, analyzed data, and drafted the manuscript; TEG was responsible for method development; DC performed method validation, SMA supervised and acquired funding; all authors reviewed and edited the manuscript.

## Data accessibility

All data supporting the findings of this manuscript are contained within the manuscript and supporting information. Raw data are available upon request from David Chakravorty.

## Abbreviations

BSA: bovine serum albumin
BODIPY: boron-dipyrromethene
GAP: GTPase activating protein
GDP: guanosine diphosphate
GEF: guanine nucleotide exchange factor
GNAI1: Human guanine nucleotide-binding protein G(i) subunit alpha-1
GNAO1: Human guanine nucleotide-binding protein G(o) subunit alpha
GPA1: Arabidopsis guanine nucleotide-binding protein alpha-1 subunit
GPCR: G protein-coupled receptor
GTP: Guanosine triphosphate
GST: glutathione S-transferase
HSLB: high salt Luria-Bertani
RGS: regulator of G protein signaling
RLK: receptor-like kinase

## Supporting Information

**Figure S1.** Comparison of GPA1 purification methods. **A.** Gel illustrating the purity of StrepII-GPA1 in our protein preparations, with the commonly co-purified ∼70 kDa DnaK band. **B.** Proteins were purified in parallel for StrepII-GPA1, His-GPA1 and GST-GPA1 (marked by *) before His-GPA1 and GST-GPA1 proteins underwent buffer exchange into “EB base”. Proteins were separated by SDS-PAGE for quantification of yield and qualitative assessment of purity. **C.** Comparison of the initial binding rates (note the x-axis units) of the StrepII-/His-/GST-GPA1 samples in Figure 1B when assayed with BODIPY-GTPγS. **D-E.** StrepII-GPA1^S52N^ does not display any binding activity when assayed with **D.** BODIPY-GTP or **E.** BODIPY-GTPγS. **F.** Traces of individual replicates averaged to calculate the StrepII-GPA1 data presented in panel **E**. Activity declined in order of the timing with which wells were assayed (rep 1→rep 2→rep 3), which corresponds to waiting time in the plate reader. **G.** Example of StrepII-GPA1 BODIPY-GTPγS binding assayed in well mode with tight error bars (compare with Fig. S1C). The assay presented in panel **C** was performed once as a control for the data in Figure 1B. The assays presented in panels **D** and **E** were performed eight times and the assays in panel **G** five times. All experiments were performed using independently purified proteins with similar results. To generate the depicted fluorescence intensity values, the raw data values of three technical replicates for each protein were baseline-corrected using the mean replicate values for the BSA negative control and averaged (except for panel **F**, which directly presents the background subtracted replicates), therefore the BSA traces appear near y=0. The data are graphically presented as the mean ± SEM, and values below zero are not plotted for panels **C** and **G**.

**Figure S2.** GDP competition with BODIPY-GTPγS and BODIPY-GTP. **A-B.** StrepII-GPA1 was purified in the presence or absence of 10 µM GDP in the binding and elution buffers. The StrepII-GPA1 sample supplemented with GDP was eluted (2.79 µM StrepII-GPA1) in the presence of 10 µM GDP. Upon dilution of StrepII-GPA1 to 100 nM final assay concentration, GDP remained in molar excess at 358 nM and competitively suppressed binding to **A.** BODIPY-GTPγS (50 nM) or **B.** BODIPY-GTP (50 nM). **C.** BODIPY-GTP binding and hydrolysis curves of StrepII-GPA1 supplemented with no GDP or 10 µM GDP and stored for 30 minutes on ice or incubated at the assay temperature of 25 °C (+incubate) prior to assay initiation. **D.** BODIPY-GTP binding and hydrolysis curves of GPA1 either freshly prepared or subjected to overnight storage at 4 °C. For the assay depicted in panel **D** the detector gain was set to 70, as opposed to 90-100 for other assays, hence the lower fluorescence intensity values. The assays presented in panels **A**, **B**, **C** and **D** were each performed twice. All replicates were performed using independently purified proteins with similar results. To generate the depicted fluorescence intensity values, the raw data values of three technical replicates for each protein were baseline-corrected using the mean replicate values for the BSA negative control and averaged, therefore the BSA traces appear near the x-axes. The data are graphically presented as the mean ± SEM and values below zero are not plotted.

**Figure S3.** Activity of StrepII-GPA1. **A.** Peak fluorescence values of BODIPY-GTP curves of StrepII-GPA1 supplemented with the indicated concentrations of the cryoprotectants glycerol or sucrose and stored at −80 °C for 3 weeks. **B.** Comparison of BODIPY-GTP binding and hydrolysis activities of StrepII-GPA1 stored at −80 °C for 3 weeks with either 10% glycerol or 8.33% sucrose added to the elution fraction (which contains 5% glycerol) as a cryoprotectant. For the assay depicted in panel **B** the detector gain was set to 70, as opposed to 90-100 for other assays, hence the lower fluorescence intensity values. The assay presented in panel **A** was performed once, with three technical replicates, while the assay presented in panel **B** was performed twice, using independently purified proteins with similar results. To generate the depicted fluorescence intensity values, the raw data values of three technical replicates for each protein were baseline-corrected using the mean replicate values for the BSA negative control and averaged, therefore the BSA traces appear near the x-axes in panel **B**. The data are graphically presented as the mean ± SEM and values below zero are not plotted.

**Figure S4.** Control data for GNAI1 codon harmonization, and buffer reagent choices. **A.** DNA sequence of the codon harmonized GNAI1ch synthesized clone. **B.** Alignment of GNA1wt (native) and GNAI1ch protein sequences, generated with Clustal Omega. **C.** SDS-PAGE illustrates the similar relative yield and purity of Strep-tag purified GNAI1wt and GNAI1ch proteins. **D-E.** Similar activities of StrepII-GNAI1wt vs. StrepII-GNAI1ch (250 nM each) for **D.** BODIPY-GTP binding and hydrolysis, and **E.** BODIPY-GTPγS binding. Note the 20-minute x-axis scale. **F.** Comparison of the BODIPY-GTP binding and hydrolysis activities of 100 nM StrepII-GPA1 purified and assayed in buffers prepared with standard (Std.) grade reagents or trace metal free (TMF) grade reagents. **G-H.** Mg^2+^ dependency for BODIPY-GTPγS binding by **G.** 200 nM GNAI1 vs. 200 nM GNAI1^S47N^ or **H.** 100 nM GNAO1 vs. 100 nM GNAO1^S47N^. The assays presented in panels **D**, **E** and **F** were performed once and the assays in panels **G** and **H** were performed twice using independently purified proteins with similar results. To generate the depicted fluorescence intensity values, the raw data values of three (panels **G** and **H**) or four (panels **D**-**F**) technical replicates for each protein were baseline-corrected using the mean replicate values for the BSA negative control and averaged, therefore the BSA traces appear near the x-axes. The data are graphically presented as the mean ± SEM and values below zero are not plotted.

**Figure S5. A-B.** Structural alignments of empirically derived **A.** GPA1-GNAO1 and **B.** GPA1-GNAI1 structures. PDB structure 2XTZ chain A (GPA1 – blue) was aligned in PyMol with 3C7K chain A (GNAO1 – green) or 1GIA chain A (GNAI1 – orange). The nucleotide (yellow for GPA1/light and dark blue for GNAO1 and GNAI1, respectively) is located in the binding pocket within the interdomain cleft, which is flanked by the Ras domain (upper domain) and helical domain (lower domain) in both panels. **C.** SYPRO Orange protein unfolding assay in the presence or absence of GTPγS for 600 nM StrepII-GNAI1 or StrepII-GNAO1. Note the reduced magnitude of fluorescence intensity values compared to StrepII-GPA1 in Figures 2A and 2B. The initial decrease in SYPRO Orange signal may result from temperature equilibration, as reactions were loaded immediately following sample preparation on ice. **D.** SYPRO Orange assay of single wells for rapid setup demonstrates that unfolding of GPA1 (600 nM) *in vitro* when not provided with excess nucleotide is almost immediate. **E.** An example of a GNAO1^GPA1hel^ SYPRO Orange protein unfolding assay in which the rapid signal variation between timepoints displayed in Figures 6C and 6D was not observed, indicating that this variation is not an inherent property of the protein. (Note the same y-axis scale was used in Figures 6C, 6D and S5E.) The assays presented in panels **C**, **D** and **E** were performed once, as additional evidence to complement main text Figures 2-5, 6C, and 6C-D, respectively. The data in panels **C** and **E** are graphically presented as the mean ± SEM of two technical replicates, while the data in panel **D** represent single measurements.

**Table S1.** Sequences of primers used in this study.

**Table S2.** Summary of calculated binding (BODIPY-GTPγS), binding and hydrolysis (BODIPY-GTP), and signal decay (SYPRO Orange) rates calculated in curve fitting analyses performed on the traces found within this manuscript.

**Table S3.** Yield comparison of StrepII-GPA1, His-GPA1 and GST-GPA1 when purified side-by-side on five different days. Yields are presented as nmol (μg).

**Table S4.** Maximal amplitudes of background-corrected BODIPY-GTP and BODIPY-GTPγS fluorescence signals derived from the data presented in this manuscript.

**Table S5.** Previously reported GPA1 GTP association rates in the literature.

**Table S6.** Clinvar data associated with equivalent sites to GNAI1^S47^/GNAO1^S47^ and GNAI1^Q204^/GNAO1^Q205^ of Gα subunits.

**Table S7.** COSMIC data associated with equivalent sites to GNAI1^S47^/GNAO1^S47^ and GNAI1^Q204^/GNAO1^Q205^ of Gα subunits.

