## Supplemental information for "Influence of expression and purification protocols on Gα biochemical activity: kinetics of plant and mammalian G protein cycles"

This file includes:
Figure S1
Figure S2
Figure S3
Figure S4
Figure S5
Table S1
Table S2
Table S3
Table S4
Table S5
Table S6
Table S7

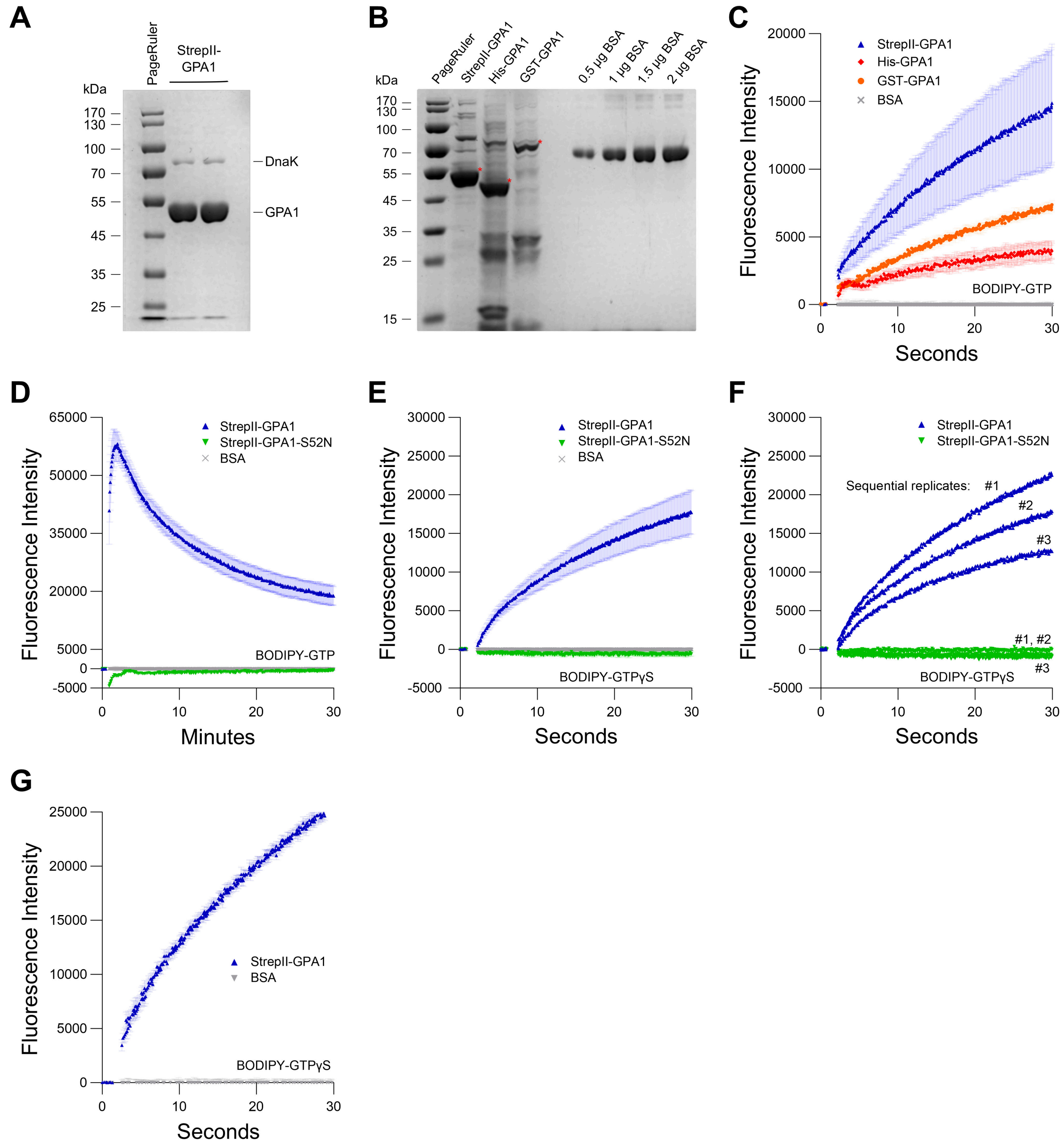


**Figure S1.** Comparison of GPA1 purification methods. **A.** Gel illustrating the purity of StrepII-GPA1 in our protein preparations, with the commonly co-purified ~70 kDa DnaK band. **B.** Proteins were purified in parallel for StrepII-GPA1, His-GPA1 and GST-GPA1 (marked by *) before His-GPA1 and GST-GPA1 proteins underwent buffer exchange into “EB base”. Proteins were separated by SDS-PAGE for quantification of yield and qualitative assessment of purity. **C.** Comparison of the initial binding rates (note the x-axis units) of the StrepII-/His-/GST-GPA1 samples in Figure 1B when assayed with BODIPY-GTPγS. **D-E.** StrepII-GPA1$^{S52N}$ does not display any binding activity when assayed with **D.** BODIPY-GTP or **E.** BODIPY-GTPγS. **F.** Traces of individual replicates averaged to calculate the StrepII-GPA1 data presented in panel **E**. Activity declined in order of the timing with which wells were assayed (rep 1→rep 2→rep 3), which corresponds to waiting time in the plate reader. **G.** Example of StrepII-GPA1 BODIPY-GTPγS binding assayed in well mode with tight error bars (compare with Fig. S1C). The assay presented in panel **C** was performed once as a control for the data in Figure 1B. The assays presented in panels **D** and **E** were performed eight times and the assays in panel **G** five times. All experiments were performed using independently purified proteins with similar results. To generate the depicted fluorescence intensity values, the raw data values of three technical replicates for each protein were baseline-corrected using the mean replicate values for the BSA negative control, and averaged (except for panel **F**, which directly presents the background subtracted replicates), therefore the BSA traces appear near y=0. The data are graphically presented as the mean ± SEM, and values below zero are not plotted for panels **C** and **G**.

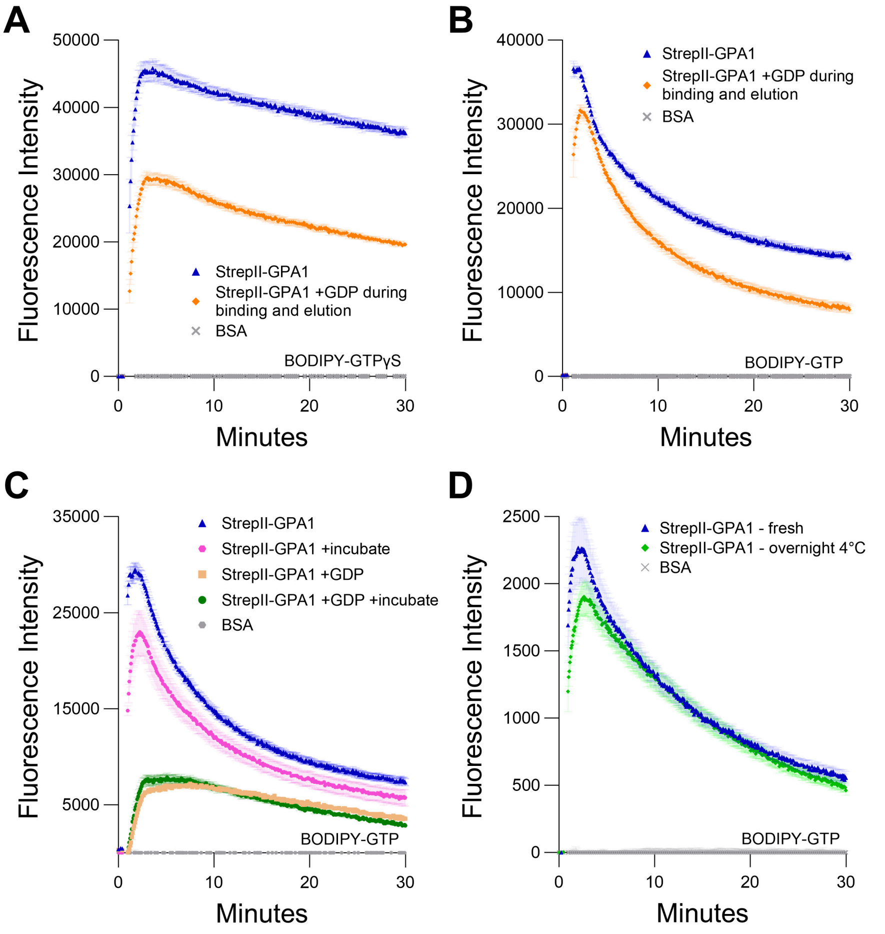


**Figure S2.** GDP competition with BODIPY-GTPγS and BODIPY-GTP. **A-B.** StrepII-GPA1 was purified in the presence or absence of 10 μM GDP in the binding and elution buffers. The StrepII-GPA1 sample supplemented with GDP was eluted (2.79 μM StrepII-GPA1) in the presence of 10 μM GDP. Upon dilution of StrepII-GPA1 to 100 nM final assay concentration, GDP remained in molar excess at 358 nM and competitively suppressed binding to **A.** BODIPY-GTPγS (50 nM) or **B.** BODIPY-GTP (50 nM). **C.** BODIPY-GTP binding and hydrolysis curves of StrepII-GPA1 supplemented with no GDP or 10 μM GDP and stored for 30 minutes on ice or incubated at the assay temperature of 25 °C (+incubate) prior to assay initiation. **D.** BODIPY-GTP binding and hydrolysis curves of GPA1 either freshly prepared or subjected to overnight storage at 4 °C. For the assay depicted in panel **D** the detector gain was set to 70, as opposed to 90-100 for other assays, hence the lower fluorescence intensity values. The assays presented in panels **A**, **B**, **C** and **D** were each performed twice. All replicates were performed using independently purified proteins with similar results. To generate the depicted fluorescence intensity values, the raw data values of three technical replicates for each protein were baseline-corrected using the mean replicate values for the BSA negative control and averaged, therefore the BSA traces appear near the x-axes. The data are graphically presented as the mean ± SEM and values below zero are not plotted.

**A**

| Cryoprotectant added | Peak BODIPY-GTP fluorescence ±SEM |
|---|---|
| None | 46510±1992 |
| 5% glycerol | 48201±1624 |
| 10% glycerol | **49779±1370** |
| 15% glycerol | 48372±903 |
| 20% glycerol | 48595±1168 |
| 25% glycerol | 48845±893 |
| 5% sucrose | 49907±135 |
| 8.3% sucrose | **51597±242** |
| 10% sucrose | 50026±747 |
| 15% sucrose | 48236±2831 |
| 20% sucrose | 46618±3872 |
| 25% sucrose | 44786±4538 |

**B**

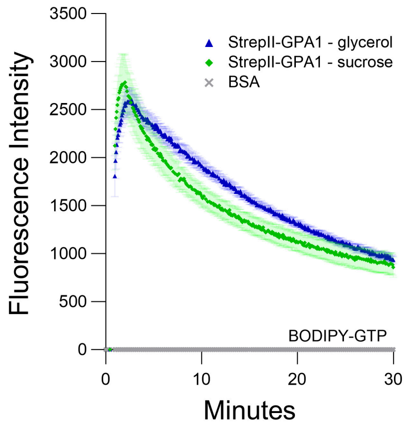


**Figure S3.** Activity of StrepII-GPA1. **A.** Peak fluorescence values of BODIPY-GTP curves of StrepII-GPA1 supplemented with the indicated concentrations of the cryoprotectants glycerol or sucrose and stored at -80 °C for 3 weeks. **B.** Comparison of BODIPY-GTP binding and hydrolysis activities of StrepII-GPA1 stored at -80 °C for 3 weeks with either 10% glycerol or 8.33% sucrose added to the elution fraction (which contains 5% glycerol) as a cryoprotectant. For the assay depicted in panel **B** the detector gain was set to 70, as opposed to 90-100 for other assays, hence the lower fluorescence intensity values. The assay presented in panel **A** was performed once, with three technical replicates, while the assay presented in panel **B** was performed twice, using independently purified proteins with similar results. To generate the depicted fluorescence intensity values, the raw data values of three technical replicates for each protein were baseline-corrected using the mean replicate values for the BSA negative control and averaged, therefore the BSA traces appear near the x-axes in panel **B**. The data are graphically presented as the mean ± SEM and values below zero are not plotted.

**A**

```
>GNAI1ch
ATGGGCTGCACCCTGTCTGCCGAAGATAAAGCGGCAGTTGAACGCTCCAAAATGATCGACCGCAACCTGCGTGAAGACGGTGAAAAAGCCGCGCGTGAAGTTAAACTGCTCCTGCTCGGTGC
AGGCGAATCTGGCAAATCCACCATCGTTAAACAGATGAAAATCATCCACGAAGCCGGTTATTCTGAAGAGGAATGTAAACAGTACAAAGCAGTGGTTTACAGCAACACCATCCAGTCTATTA
TCGCTATCATTCGTGCGATGGGTCGTCTGAAAATCGACTTTGGTGACAGCGCCCGTGCGGATGACGCACGCCAGCTGTTTGTGCTGGCAGGTGCAGCGGAGGAAGGCTTTATGACTGCAGAA
CTGGCCGGTGTTATCAAACGTCTGTGGAAAGATTCTGGTGTACAGGCCTGTTTCAACCGTTCCCGTGAATACCAGCTGAACGATTCTGCAGCGTACTATCTGAACGACCTGGACCGTATCGC
TCAGCCGAACTACATCCCGACTCAGCAGGATGTTCTGCGTACTCGTGTGAAAACTACCGGTATTGTTGAAACCCACTTTACTTTCAAAGATCTGCACTTTAAAATGTTTGATGTGGGTGGCC
AGCGTTCTGAACGTAAGAAATGGATTCACTGCTTCGAAGGTGTGACCGCGATTATCTTCTGTGTAGCACTGTCTGACTACGACCTGGTTCTGGCCGAAGATGAGGAAATGAACCGTATGCAC
GAAAGCATGAAACTGTTTGACAGCATCTGTAATAACAAATGGTTTACCGATACCAGCATTATCCTGTTTCTGAACAAAAAAGATCTGTTTGAGGAAAAAATCAAGAAAAGCCCGCTGACCAT
CTGCTACCCGGAATATGCGGGTTCTAACACCTATGAAGAGGCGGCTGCATATATTCAGTGTCAGTTTGAAGACCTGAACAAACGTAAAGACACCAAAGAAATCTACACCCACTTCACCTGTG
CCACCGATACTAAAAACGTGCAGTTTGTTTTTGATGCTGTAACCGATGTGATCATTAAAAATAACCTGAAGGATTGCGGTCTGTTCTAA
```

**B**

```
GNAI1wt      MGCTLSAEDKAAVERSKMIDRNLREDGEKAAREVKLLLLGAGESGKSTIVKQMKIIHEAG   60
GNAI1ch      MGCTLSAEDKAAVERSKMIDRNLREDGEKAAREVKLLLLGAGESGKSTIVKQMKIIHEAG   60
             ************************************************************

GNAI1wt      YSEEECKQYKAVVYSNTIQSIIAIIRAMGRLKIDFGDSARADDARQLFVLAGAAEEGFMT   120
GNAI1ch      YSEEECKQYKAVVYSNTIQSIIAIIRAMGRLKIDFGDSARADDARQLFVLAGAAEEGFMT   120
             ************************************************************

GNAI1wt      AELAGVIKRLWKDSGVQACFNRSREYQLNDSAAYYLNDLDRIAQPNYIPTQQDVLRTRVK   180
GNAI1ch      AELAGVIKRLWKDSGVQACFNRSREYQLNDSAAYYLNDLDRIAQPNYIPTQQDVLRTRVK   180
             ************************************************************

GNAI1wt      TTGIVETHFTFKDLHFKMFDVGGQRSERKKWIHCFEGVTAIIFCVALSDYDLVLAEDEEM   240
GNAI1ch      TTGIVETHFTFKDLHFKMFDVGGQRSERKKWIHCFEGVTAIIFCVALSDYDLVLAEDEEM   240
             ************************************************************

GNAI1wt      NRMHESMKLFDSICNNKWFTDTSIILFLNKKDLFEEKIKKSPLTICYPEYAGSNTYEEAA   300
GNAI1ch      NRMHESMKLFDSICNNKWFTDTSIILFLNKKDLFEEKIKKSPLTICYPEYAGSNTYEEAA   300
             ************************************************************

GNAI1wt      AYIQCQFEDLNKRKDTKEIYTHFTCATDTKNVQFVFDAVTDVIIKNNLKDCGLF*   354
GNAI1ch      AYIQCQFEDLNKRKDTKEIYTHFTCATDTKNVQFVFDAVTDVIIKNNLKDCGLF*   354
             *******************************************************
```

**C**

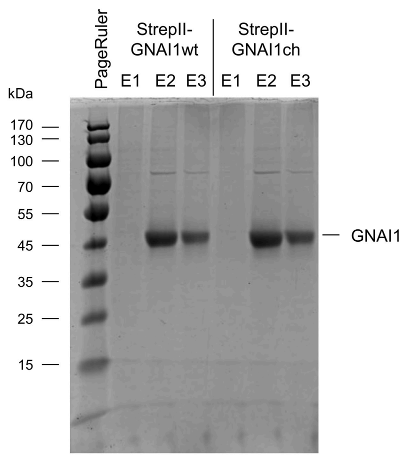


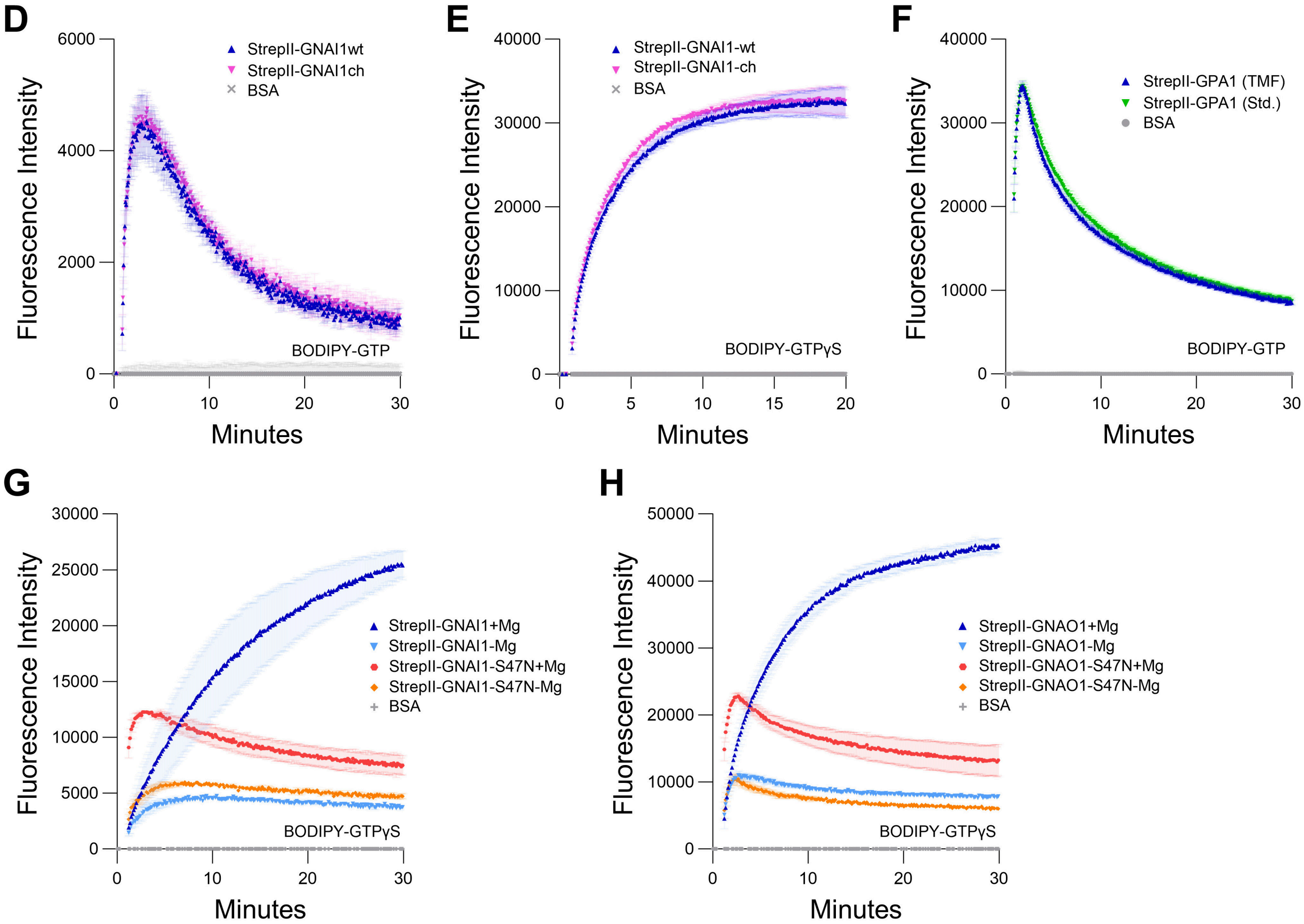


**Figure S4.** Control data for GNAI1 codon harmonization, and buffer reagent choices. **A.** DNA sequence of the codon harmonized GNAI1ch synthesized clone. **B.** Alignment of GNA1wt (native) and GNAI1ch protein sequences, generated with Clustal Omega. **C.** SDS-PAGE illustrates the similar relative yield and purity of Strep-tag purified GNAI1wt and GNAI1ch proteins. **D-E.** Similar activities of StrepII-GNAI1wt vs. StrepII-GNAI1ch (250 nM each) for **D.** BODIPY-GTP binding and hydrolysis, and **E.** BODIPY-GTPγS binding. Note the 20-minute x-axis scale. **F.** Comparison of the BODIPY-GTP binding and hydrolysis activities of 100 nM StrepII-GPA1 purified and assayed in buffers prepared with standard (Std.) grade reagents or trace metal free (TMF) grade reagents. **G-H.** $Mg^{2+}$ dependency for BODIPY-GTPγS binding by **G.** 200 nM GNAI1 vs. 200 nM $GNAI1^{S47N}$ or **H.** 100 nM GNAO1 vs. 100 nM $GNAO1^{S47N}$. The assays presented in panels **D**, **E** and **F** were performed once and the assays in panels **G** and **H** were performed twice using independently purified proteins with similar results. To generate the depicted fluorescence intensity values, the raw data values of three (panels **G** and **H**) or four (panels **D-F**) technical replicates for each protein were baseline-corrected using the mean replicate values for the BSA negative control and averaged, therefore the BSA traces appear near the x-axes. The data are graphically presented as the mean ± SEM and values below zero are not plotted.

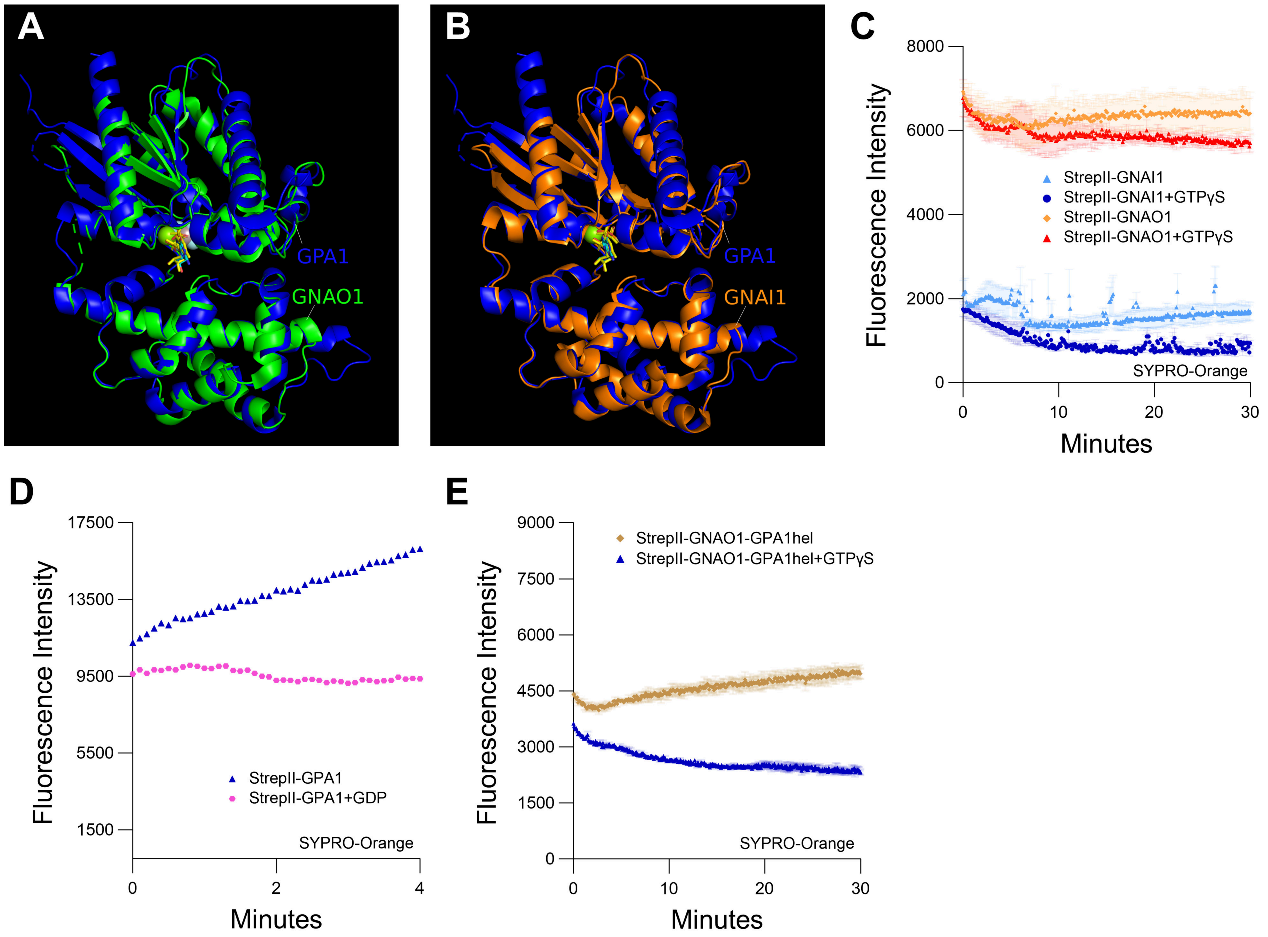


**Figure S5. A-B.** Structural alignments of empirically derived **A.** GPA1-GNAO1 and **B.** GPA1-GNAI1 structures. PDB structure 2XTZ chain A (GPA1 – blue) was aligned in PyMol with 3C7K chain A (GNAO1 – green) or 1GIA chain A (GNAI1 – orange). The nucleotide (yellow for GPA1/light and dark blue for GNAO1 and GNAI1, respectively) is located in the binding pocket within the interdomain cleft, which is flanked by the Ras domain (upper domain) and helical domain (lower domain) in both panels. **C.** SYPRO Orange protein unfolding assay in the presence or absence of GTPγS for 600 nM StrepII-GNAI1 or StrepII-GNAO1. Note the reduced magnitude of fluorescence intensity values compared to StrepII-GPA1 in Figures 2A and 2B. The initial decrease in SYPRO Orange signal may result from temperature equilibration, as reactions were loaded immediately following sample preparation on ice. **D.** SYPRO Orange assay of single wells for rapid setup demonstrates that unfolding of GPA1 (600 nM) in vitro when not provided with excess nucleotide is almost immediate. **E.** An example of a $GNAO1^{GPA1hel}$ SYPRO Orange protein unfolding assay in which the rapid signal variation between timepoints displayed in Figures 6C and 6D was not observed, indicating that this variation is not an inherent property of the protein. (Note the same y-axis scale was used in Figures 6C, 6D and S5E.) The assays presented in panels **C**, **D** and **E** were performed once, as additional evidence to complement main text Figures 2-5, 6C, and 6C-D, respectively. The data in panels **C** and **E** are graphically presented as the mean ± SEM of two technical replicates, while the data in panel **D** represent single measurements.

**Table S1.** Sequences of primers used in this study.

| Primer | Sequence | Use |
|---|---|---|
| GPA1f-NcoI | GGGCCATGGGCTTACTCTGCAG | Cloning *GPA1* into pSTTa |
| GPA1r-BspEI | GGGTCCGGATCATAAAAGGCCAGCCTCCAG | |
| GPA1f-BspHI | GCGTCATGATGGGCTTACTCTGCAG | Cloning the *GPA1*$^{GNAO1hel}$ construct into pSTTa |
| GNAI1f-NcoI | GCGCCATGGGCTGCACCCTGTC | Cloning codon harmonized *GNAI1* into pSTTa |
| GNAI1r-BspEI | GGGTCCGGATTAGAACAGACCGCAATCCTTC | |
| GNAI1f-wt-NcoI | GCACCATGGGCTGCACGCTGAG | Cloning the native (wild-type) *GNAI1* into pSTTa |
| GPA1-37f-NcoI | GCGCCATGGGTCATATTCGGAAGCTTTTGCTAC | |
| GNAO1f-BspHI | GCGTCATGATGGGATGTACTCTGAGCG | Cloning *GNAO1* into pSTTa |
| GNAO1r | TCAGTACAAGCCGCAGCCC | |
| GPA1S52NF | CTGGAAAAAAATACAATTTTTAAGC | S52N mutagenesis of *GPA1* |
| GPA1S52NR | GCTTAAAAATTGTATTTTTTCCAG | |
| GNAI1-S47N-rf | GGCAAAAACACCATCGTTAAACAGATGAAAATCATCC | S47N mutagenesis of *GNAI1* |
| GNAI1-S47N-rr | ACGATGGTGTTTTTGCCAGATTCGCCTGC | |
| GNAI1-Q204L-rf | GGGTGGCCTGCGTTCTGAACGTAAGAAATGG | Q204L mutagenesis of *GNAI1* |
| GNAI1-Q204L-rr | CAGAACGCAGGCCACCCACATCAAACAT | |
| GNAO1-S47N-rf | AGGAAAAAAACACCATTGTGAAGCAGATGAAG | S47N mutagenesis of *GNAO1* |
| GNAO1-S47N-rr | CAATGGTGTTTTTTCCTGATTCTCCAGC | |
| GNAO1-Q205L-rf | CGGAGGCCTGCGATCTGAACGCAAGAAG | Q205L mutagenesis of *GNAO1* |
| GNAO1-Q205L-rr | CAGATCGCAGGCCTCCGACGTCAAACAG | |
| GPA1-GNAO1hel-or | GTACTGTTTCACGTCTTCTCCATCAAATCCCGTTTGGAATAGAAG | Reverse primer of the N-terminal fragment of the *GPA1* Ras domain (for overlap extension PCR) |
| GPA1-GNAO1hel-of | GAGCAGGACATCCTCCGAGCAAGAGTTCGCACAACTG | Forward primer of the C-terminal fragment of the *GPA1* Ras domain (for overlap extension PCR) |
| GPA1hel-f2 | GAAGGAGAACTAAAGAGCTATGTTC | Forward primer of the *GPA1* helical domain (for overlap extension PCR) |
| GPA1hel-r | ATAAAGTACATCCTCCTTAGTTGG | Reverse primer of the *GPA1* helical domain (for overlap extension PCR) |
| GNAO1-GPA1hel-or | GAACATAGCTCTTTAGTTCTCCTTCGGAGAAGCCATCTTCATGGATG | Reverse primer of the N-terminal fragment of the |

| | | |
|---|---|---|
| | | *GNAO1* Ras domain (for overlap extension PCR) |
| GNAO1-GPA1hel-of | CCAACTAAGGAGGATGTACTTTATACCAGGGTCAAAACCACTG | Forward primer of the C-terminal fragment of the *GNAO1* Ras domain (for overlap extension PCR) |
| GNAO1hel-f | GGAGAAGACGTGAAACAGTACAAG | Forward primer of the *GNAO1* helical domain (for overlap extension PCR) |
| GNAO1hel-r | TCGGAGGATGTCCTGCTC | Reverse primer of the *GNAO1* helical domain (for overlap extension PCR) |
| GPA1f | ATGGGCTTACTCTGCAGTA | Cloning *GPA1* into pCR8 |
| GPA1rs | TCATAAAAGGCCAGCCTCC | |
| GNAI1f | ATGGGCTGCACCCTGTCTG | Cloning codon harmonized *GNAI1* into pCR8 |
| GNAI1r | TTAGAACAGACCGCAATCCTTCAG | |
| GNAO1f | ATGGGATGTACTCTGAGCG | Cloning *GNAO1* into pCR8 |
| GNAO1r | TCAGTACAAGCCGCAGCCC | |
| RGS1 NcoI 763F | GGATCCATGGGCTCGAGTAACCAACCTCTTCTTTCAC | Cloning the region corresponding to the C-terminal cytosolic RGS domain of *RGS1* into pSTTa |
| RGS1 BspEI 1355Rs | TCAAGCTCCGGATTAACCGGGACTACTGCATCTGGAAC | |

**Table S2.** Summary of calculated binding (BODIPY-GTPγS), binding and hydrolysis (BODIPY-GTP), and signal decay (SYPRO Orange) rates calculated in curve fitting analyses performed on the traces found within this manuscript.

| | | Observed association | | | Observed decay | | | |
|---|---|---|---|---|---|---|---|---|
| | | k ($min^{-1}$) | $R^2$ | k as % | k ($min^{-1}$) | $R^2$ | k as % | |
| **Fig. 1B** | StrepII-GPA1 | 3.139 | 0.980 | 100.0 | 0.09426 | 0.9988 | 100 | The average observed BODIPY-GTP |
| BODIPY-GTP | His-GPA1 | 1.216 | 0.970 | 38.7 | 0.01391 | 0.9975 | 14.8 | association rate for Strep-GPA1 over |
| | GST-GPA1 | 2.415 | 0.991 | 76.9 | 0.0575 | 0.9984 | 61 | seven figures = 3.192 ± 0.120 (SEM) $min^{-1}$ |
| **Fig. 2D** | GTPγS T=30 min | N.D. | N.D. | N.D. | 0.077 | 0.988 | 100 | |
| SYPRO-Orange | GTPγS T=20 min | N.D. | N.D. | N.D. | 0.113 | 0.984 | 146.9 | |
| | GTPγS T=15 min | N.D. | N.D. | N.D. | 0.185 | 0.989 | 239.6 | |
| | GTPγS T=10 min | N.D. | N.D. | N.D. | 0.291 | 0.975 | 378.0 | |
| | GTPγS T=5 min | N.D. | N.D. | N.D. | 0.682 | 0.964 | 885.0 | |
| **Fig. 4A** | StrepII-GPA1 | 3.156 | 0.955 | 100.0 | 0.1395 | 0.9908 | 100.0 | |
| BODIPY-GTP | StrepII-GNAO1 | 1.065 | 0.999 | 33.7 | 0.1512 | 0.9975 | 108.4 | |
| | StrepII-GNAI1 | 0.844 | 0.707 | 26.7 | 0.1094 | 0.9602 | 78.4 | |
| **Fig. 4C** | StrepII-GNAO1 | 0.671 | 0.968 | 100.0 | 0.09089 | 0.9982 | 100.0 | |
| BODIPY-GTP | His-GNAO1 | 1.220 | 0.958 | 181.8 | 0.2629 | 0.9982 | 289.3 | |
| **Fig. 4E** | StrepII-GPA1 | 3.251 | 0.823 | 100.0 | 0.1127 | 0.9975 | 100.0 | |
| BODIPY-GTP | His-GNAO1 | 0.801 | 0.968 | 24.6 | 0.2016 | 0.9976 | 178.9 | |
| **Fig. 4F** | StrepII-GPA1 | 1.442 | 0.977 | 100.0 | N.D. | N.D. | N.D. | |
| BODIPY-GTPγS | His-GNAO1 | 0.206 | 0.999 | 14.3 | N.D. | N.D. | N.D. | |
| **Fig. 5A** | **StrepII-GPA1** | | | | | | | |
| BODIPY-GTPγS | 1.2 μM | 3.531 | 0.9278 | N.D. | N.D. | N.D. | N.D. | Our curve fitting approach for BODIPY-GTP is |
| | 800 nM | 3.364 | 0.9326 | N.D. | N.D. | N.D. | N.D. | validated by the protein saturation binding |
| | 400 nM | 2.894 | 0.9209 | N.D. | N.D. | N.D. | N.D. | kinetic values obtained using BODIPY-GTPγS. |
| | 200 nM | 1.946 | 0.7887 | N.D. | N.D. | N.D. | N.D. | |
| | 100 nM | 1.592 | 0.253 | N.D. | N.D. | N.D. | N.D. | |
| **Fig. 5B** | **His-GNAO1** | | | | | | | |
| BODIPY-GTPγS | 1.2 μM | 1.805 | 0.973 | N.D. | N.D. | N.D. | N.D. | |
| | 800 nM | 1.461 | 0.9667 | N.D. | N.D. | N.D. | N.D. | |
| | 400 nM | 0.9524 | 0.9737 | N.D. | N.D. | N.D. | N.D. | |
| | 200 nM | 0.4934 | 0.9916 | N.D. | N.D. | N.D. | N.D. | |
| | 100 nM | 0.2798 | 0.9978 | N.D. | N.D. | N.D. | N.D. | |
| **Fig. 6A** | StrepII-GPA1 | 1.126 | 0.965 | 100.0 | N.D. | N.D. | N.D. | |
| BODIPY-GTPγS | StrepII-GPA1-GNAO1hel | 0.533 | 0.996 | 47.4 | N.D. | N.D. | N.D. | |
| | StrepII-GNAO1 | 0.134 | 0.999 | 11.9 | N.D. | N.D. | N.D. | |
| | StrepII-GNAO1-GPA1hel | 1.608 | 0.955 | 142.8 | N.D. | N.D. | N.D. | |
| **Fig. 6B** | StrepII-GPA1 | 3.088 | 0.972 | 100.0 | 0.144 | 0.992 | 100 | |
| BODIPY-GTP | StrepII-GPA1-GNAO1hel | 0.867 | 0.911 | 28.1 | 0.116 | 0.999 | 80.7 | |
| | StrepII-GNAO1 | 0.806 | 0.900 | 26.1 | 0.117 | 0.996 | 81.4 | |
| | StrepII-GNAO1-GPA1hel | 2.651 | 0.952 | 85.8 | 0.505 | 0.993 | 351.5 | |
| **Fig. S2A** | StrepII-GPA1 | 1.478 | 0.973 | 100.0 | N.D. | N.D. | N.D. | |
| BODIPY-GTPγS | StrepII-GPA1 with GDP-binding/elution | 1.137 | 0.964 | 76.9 | N.D. | N.D. | N.D. | |
| **Fig. S2B** | StrepII-GPA1 | 3.356 | 0.945 | 100.0 | 0.137 | 0.991 | 100.0 | |
| BODIPY-GTP | StrepII-GPA1 with GDP-binding/elution | 2.272 | 0.9803 | 67.7 | 0.136 | 0.997 | 100.5 | |
| **Fig. S2C** | StrepII-GPA1 | 3.706 | 0.965 | 100.0 | 0.136 | 0.997 | 100.0 | |
| BODIPY-GTP | StrepII-GPA1+GDP | 0.718 | 0.923 | 19.4 | 0.018 | 0.993 | 12.9 | |
| | StrepII-GPA1+incubate | 2.123 | 0.983 | 57.3 | 0.121 | 0.997 | 89.3 | |
| | StrepII-GPA1+GDP+incubate | 0.819 | 0.909 | 22.1 | 0.034 | 0.997 | 25.2 | |
| **Fig. S2D** | StrepII-GPA1 - fresh | 2.648 | 0.989 | 100.0 | 0.091 | 0.998 | 100.0 | |
| BODIPY-GTP | StrepII-GPA1 - overnight 4°C | 1.814 | 0.994 | 68.5 | 0.055 | 0.999 | 60.1 | |
| **Fig. S3B** | StrepII-GPA1 - sucrose | 3.032 | 0.9786 | 100 | 0.104 | 0.9958 | 100 | |
| BODIPY-GTP | StrepII-GPA1 - glycerol | 2.339 | 0.9896 | 77.1 | 0.047 | 0.9993 | 44.8 | |

**Table S3.** Yield comparison of StrepII-GPA1, His-GPA1 and GST-GPA1 when purified side-by-side on five different days. Yields are presented as nmol (μg).

| | Set 1 | Set 2 | Set 3 | Set 4 | Set 5 | Average |
|---|---|---|---|---|---|---|
| StrepII-GPA1 | 0.67 (33.6) | 0.74 (37.0) | 0.96 (47.9) | 1.09 (54.7) | 1.20 (60.0) | 0.93 (46.6) |
| His-GPA1 | 0.46 (22.1) | 0.56 (25.9) | 0.37 (17.4) | 0.53 (25.3) | 0.65 (31.1) | 0.51 (24.4) |
| GST-GPA1 | 0.11 (7.9) | 0.10 (6.9) | 0.13 (9.5) | 0.14 (10.2) | 0.11 (7.7) | 0.12 (8.4) |

**Table S4.** Maximal amplitudes of background-corrected BODIPY-GTP and BODIPY-GTPγS fluorescence signals derived from the data presented in this manuscript

| Figure | Statistic | Analyzed proteins | | | | | |
|---|---|---|---|---|---|---|---|
| | | StrepII-GPA1 | His-GPA1 | GST-GPA1 | BSA | | |
| 1B | Maximum | 43368 | 23353 | 32353 | 1E-12 | | |
| | | StrepII-GPA1 | StrepII-GPA1 + StrepII-RGS1c (1:2) | StrepII-GPA1 + StrepII-RGS1c (1:4) | | | |
| 1C | Maximum | 17868 | 15806 | 12007 | | | |
| | | StrepII-GPA1 | StrepII-GPA1+GDP | Buffer | | | |
| 2A | Maximum | 27639 | 15902 | 0 | | | |
| | | StrepII-GPA1 | StrepII-GPA1+GTPγS | Buffer | | | |
| 2B | Maximum | 21683 | 9736 | 0 | | | |
| | | StrepII-GNAI1 | StrepII-GNAI1-S47N | StrepII-GNAI1-Q204L | BSA | | |
| 3A | Maximum | 2577 | 706 | 7406 | 6E-13 | | |
| | | StrepII-GNAO1 | StrepII-GNAO1-S47N | StrepII-GNAO1-Q205L | BSA | | |
| 3B | Maximum | 14435 | 867 | 6236 | 6E-13 | | |
| | | StrepII-GNAI1 | StrepII-GNAI1-S47N | StrepII-GNAI1-Q204L | BSA | | |
| 3C | Maximum | 26196 | 12680 | 18212 | 6E-13 | | |
| | | StrepII-GNAO1 | StrepII-GNAO1-S47N | StrepII-GNAO1-Q205L | BSA | | |
| 3D | Maximum | 45862 | 23743 | 12356 | 6E-13 | | |
| | | StrepII-GPA1 | StrepII-GNAO1 | StrepII-GNAI1 | BSA | | |
| 4A | Maximum | 42897 | 19055 | 2555 | 1E-12 | | |
| | | StrepII-GNAI1 | His-GNAI1 | BSA | | | |
| 4B | Maximum | 3063 | 3942 | 1E-12 | | | |
| | | StrepII-GNAO1 | His-GNAO1 | BSA | | | |
| 4C | Maximum | 9231 | 33077 | 6E-13 | | | |
| | | StrepII-GNAO1 | His-GNAO1 | BSA | | | |
| 4D | Maximum | 24332 | 29422 | 3E-13 | | | |
| | | StrepII-GPA1 | His-GNAO1 | BSA | | | |
| 4E | Maximum | 23300 | 14629 | 6E-13 | | | |
| | | StrepII-GPA1 | His-GNAO1 | BSA | | | |
| 4F | Maximum | 21945 | 51177 | 1E-12 | | | 5A and 5B share the BSA sample |
| | | 100 nM GPA1 | 200 nM GPA1 | 400 nM GPA1 | 800 nM GPA1 | 1.2 µM GPA1 | BSA |
| 5A | Maximum | 5179 | 9592 | 12331 | 13111 | 13181 | 0 |
| | | 100 nM GNAO1 | 200 nM GNAO1 | 400 nM GNAO1 | 800 nM GNAO1 | 1.2 µM GNAO1 | BSA |
| 5B | Maximum | 10292 | 10711 | 10652 | 10613 | 10593 | 0 |
| | | StrepII-GPA1+GDP | StrepII-GPA1 | His-GNAO1+GDP | His-GNAO1 | BSA | |
| 5C | Maximum | 8223 | 23632 | -86 | 12233 | 6E-13 | |
| | | StrepII-GPA1 | StrepII-GPA1-GNAO1hel | StrepII-GNAO1 | StrepII-GNAO1-GPA1hel | BSA | |
| 6A | Maximum | 39991 | 50383 | 50320 | 64860 | 6E-13 | |
| | | StrepII-GPA1 | StrepII-GPA1-GNAO1hel | StrepII-GNAO1 | StrepII-GNAO1-GPA1hel | BSA | |
| 6B | Maximum | 32316 | 44123 | 11572 | 11059 | 6E-13 | |
| | | StrepII-GPA1 | His-GPA1 | GST-GPA1 | BSA | | |
| S1C | Maximum | 15211 | 4317 | 7662 | 0 | | |
| | | StrepII-GPA1 | StrepII-GPA1-S52N | BSA | | | |
| S1D | Maximum | 58194 | 34 | 1E-12 | | | |
| | | StrepII-GPA1 | StrepII-GPA1-S52N | BSA | | | |
| S1E | Maximum | 18506 | 109 | 1E-12 | | | |
| | | StrepII-GPA1 (Rep #1) | StrepII-GPA1 (Rep #2) | StrepII-GPA1 (Rep #3) | StrepII-GPA1-S52N (Rep #1) | StrepII-GPA1-S52N (Rep #2) | StrepII-GPA1-S52N (Rep #3) |
| S1F | Maximum | 23570 | 18638 | 13389 | 379 | 161 | 81 |
| | | StrepII-GPA1 | BSA | | | | |
| S1G | Maximum | 25763 | 2E-12 | | | | |
| | | StrepII-GPA1 | StrepII-GPA1 +GDP-binding/elution | BSA | | | |
| S2A | Maximum | 45839 | 29542 | 6E-13 | | | |
| | | StrepII-GPA1 | StrepII-GPA1 GDP-binding/elution | BSA | | | |
| S2B | Maximum | 36645 | 31655 | 6E-13 | | | |
| | | StrepII-GPA1 | StrepII-GPA1 +GDP | StrepII-GPA1 +incubate | StrepII-GPA1 +GDP +incubate | BSA | |
| S2C | Maximum | 29485 | 7212 | 23011 | 7833 | 6E-13 | |
| | | StrepII-GPA1 - fresh | StrepII-GPA1 - overnight 4°C | BSA | | | |
| S2D | Maximum | 2269 | 1904 | 0 | | | |
| | | StrepII-GPA1 - sucrose | StrepII-GPA1 - glycerol | BSA | | | |
| S3B | Maximum | 2786 | 2605 | 0 | | | |
| | | StrepII-GNAI1wt | StrepII-GNAI1ch | BSA | | | |
| S4D | Maximum | 4527 | 4756 | 0 | | | |
| | | StrepII-GNAI1-wt | StrepII-GNAI1-ch | BSA | | | |
| S4E | Maximum | 32761 | 32948 | 0 | | | |
| | | StrepII-GPA1 (TMF) | StrepII-GPA1 (Std.) | BSA | | | |
| S4F | Maximum | 34352 | 34438 | 0 | | | |
| | | StrepII-GNAI1+Mg | StrepII-GNAI1-Mg | StrepII-GNAI1-S47N+Mg | StrepII-GNAI1-S47N-Mg | BSA | |
| S4G | Maximum | 25918 | 4754 | 12264 | 5994 | 6E-13 | |
| | | StrepII-GNAO1+Mg | StrepII-GNAO1-Mg | StrepII-GNAO1-S47N+Mg | StrepII-GNAO1-S47N-Mg | BSA | |
| S4H | Maximum | 45718 | 11022 | 22797 | 10563 | 6E-13 | |
| | | StrepII-GPA1 | StrepII-GPA1+GDP | | | | |
| S5D | Maximum | 16143 | 9371 | | | | |
| | | StrepII-GNAO1-GPA1hel | StrepII-GNAO1-GPA1hel+GTPγS | | | | |
| S5E | Maximum | 5055 | 3623 | | | | |

**Table S5.** Previously reported GPA1 GTP association rates in the literature.

| **Calculated GTP association rates of GPA1** | **Method** | **Study** |
|---|---|---|
| 1.44 $min^{-1}$ | $[^{35}S]$GTPγS binding | {Johnston, 2007 #518} |
| 14.4 $min^{-1}$ | BODIPY-GTPγS binding | {Johnston, 2007 #518} |
| 1.7 $min^{-1}$ | Intrinsic tryptophan fluorescence | {Jones, 2011 #1280} |
| 3.4 $min^{-1}$ | MANT-GMPPNP binding | {Jones, 2011 #1278} |
| 4.1 $min^{-1}$ | Intrinsic tryptophan fluorescence | {Jones, 2011 #1278} |
| 5.8 $min^{-1}$ | $[^{35}S]$GTPγS binding | {Urano, 2012 #1864} |
| 3.63 $min^{-1}$ | Intrinsic tryptophan fluorescence | {Urano, 2012 #1864} |
| 2.6 $min^{-1}$ | Intrinsic tryptophan fluorescence | {Jones, 2012 #1848} |
| 0.9 $min^{-1}$ | [35S]GTPγS binding | {Jones, 2012 #1848} |

**Table S6.** Clinvar data associated with equivalent sites to $GNAI1^{S47}$/$GNAO1^{S47}$ and $GNAI1^{Q204}$/$GNAO1^{Q205}$ of Gα subunits.

| **Gene** | **Mutations to protein** | **Conditions** |
|---|---|---|
| *GNAI3* | S47R | Auriculocondylar syndrome 1 |
| *GNAI3* | S47N | Auriculocondylar syndrome 1 |
| *GNAO1* | S47N | Developmental and epileptic encephalopathy 17, early infantile epileptic encephalopathy with suppression bursts |
| *GNAT2* | S47G | Achromatopsia 4 |
| *GNAT3* | S47T | Inborn genetic diseases |
| *GNAI1* | Q204R | Neurodevelopmental disorder with hypotonia, impaired speech, and behavioral abnormalities |
| *GNAO1* | Q205P | Developmental and epileptic encephalopathy 17 |
| *GNAO1* | Q205L | Early infantile epileptic encephalopathy with suppression bursts |
| *GNAT1* | Q200E | Congenital stationary night blindness autosomal dominant 3 |
| *GNAS* | Q227K, Q227H | McCune-Albright syndrome |
| *GNAS* | Q227L | Neoplasm |
| *GNAS* | Q227R, Q227H | Pituitary adenoma 3, multiple types |
| *GNAQ* | Q209H | Familial multiple nevi flammei |
| *GNAQ* | Q209L, Q209P | Melanoma, uveal melanoma |

| | | |
|---|---|---|
| *GNAQ* | Q209R | Melanoma, abnormality of cardiovascular system morphology, Sturge-Weber syndrome |
| *GNA11* | Q209R, Q209E | Not provided |
| *GNA11* | Q209P, Q209L | Melanoma, uveal melanoma |
| *GNA14* | Q205L | Congenital tufted angioma, Kaposiform hemangioendothelioma, Pyogenic granuloma |

**Table S7.** COSMIC data associated with equivalent sites to GNAI1$^{S47}$/GNAO1$^{S47}$ and GNAI1$^{Q204}$/GNAO1$^{Q205}$ of Gα subunits.

| **Gene** | **Mutations to protein** | **Conditions** | **Samples** |
|---|---|---|---|
| *GNAI2* | S47N | Lymphoid neoplasm, large intestine carcinoma | 7 |
| *GNAT1* | S43N | Large intestine carcinoma | 1 |
| *GNAS* | S54R | Malignant melanoma | 1 |
| *GNAL* | S56I | Liver neoplasm | 1 |
| *GNA11* | S53G | Large intestine carcinoma | 3 |
| *GNA13* | S62P | Burkitt lymphoma | 3 |
| *GNA15* | S56R | Ewings sarcoma-peripheral primitive neuroectodermal tumour | 1 |
| *GNAI3* | Q204R | Cervical carcinoma, Malignant melanoma | 2 |
| *GNAI3* | Q204E | Malignant melanoma | 1 |
| *GNAS* | Q227L | Malignant melanoma, lung, pancreatic, liver and salivary gland carcinomas, pituitary and stomach adenomas, fibrous dysplasia | 47 |
| *GNAS* | Q227R | Biliary tract, small intestine, large intestine, lung and pancreatic carcinomas, pituitary, stomach, thyroid (Adenoma-nodule-goitre) and pituitary adenomas | 25 |

| | | | |
|---|---|---|---|
| *GNAS* | Q227H | Adrenal gland, lung, prostate, small intestine and thyroid carcinomas, small intestine and pituitary adenomas, Adenoma-nodule-goitre (thyroid), Haematopoietic and pancreatic neoplasms, Pseudomyxoma peritonei | 21 |
| *GNAS* | Q227K | Lung and thyroid carcinomas, sex cord-stromal tumour (ovary), pituitary adenoma | 5 |
| *GNAS* | Q227E | Adrenal cortical adenoma, pancreatic carcinoma, soft tissue myxoma, adenoma-nodule-goitre (thyroid) | 5 |
| *GNAS* | Q227P | Pancreatic neoplasm | 1 |
| *GNAQ* | Q209L | Uveal , skin and central nervous system malignant melanomas, uveal and meninges melanocytomas, benign melanocytic nevus, soft tissue haemangioma | 406 |
| *GNAQ* | Q209P | Uveal and skin malignant melanomas, central nervous system glioma, germ cell tumour of the testis, meninges melanocytoma, benign melanocytic nevus, soft tissue haemangioma | 378 |
| *GNAQ* | Q209H | Uveal and skin malignant melanomas, benign melanocytic nevus, soft tissue haemangioma | 42 |
| *GNAQ* | Q209R | Uveal and skin malignant melanomas, benign melanocytic nevus, soft tissue haemangioma | 18 |
| *GNAQ* | Q209K | Uveal malignant melanoma | 1 |
| *GNAQ* | Q209Y | Uveal malignant melanoma | 1 |

| | | | |
|---|---|---|---|
| *GNA11* | Q209L | Uveal and skin malignant melanomas, meninges melanocytoma, benign melanocytic nevus, soft tissue haemangioma | 553 |
| *GNA11* | Q209H | Uveal and skin malignant melanomas, adrenal cortical adenoma, soft tissue haemangioma | 16 |
| *GNA11* | Q209P | Uveal malignant melanoma, meninges melanocytoma, soft tissue haemangioma, ovarian malignant potential (borderline) tumour | 10 |
| *GNA11* | Q209R | Uveal and skin malignant melanomas | 6 |
| *GNA11* | Q209V | Uveal malignant melanoma | 2 |
| *GNA11* | Q209A | Uveal malignant melanoma | 1 |
| *GNA11* | Q209K | Prostate carcinoma | 1 |
| *GNA13* | Q226E | Skin malignant melanomas | 1 |
